# The testis-specific transcription factor TCFL5 responds to A-MYB to elaborate the male meiotic program in placental mammals

**DOI:** 10.1101/2021.04.04.438419

**Authors:** Deniz M. Özata, Tianxiong Yu, Katharine Cecchini, Haiwei Mou, Amena Arif, Cansu Colpan, Adriano Biasini, Ildar Gaitendinov, Dirk G. de Rooij, Zhiping Weng, Phillip D. Zamore

## Abstract

In male mice, the transcription factor (TF) A-MYB initiates reprogramming of gene expression after spermatogonia enter meiosis. We report that A-MYB activates *Tcfl5*, a testis-specific TF first produced in pachytene spermatocytes. Subsequently, A-MYB and TCFL5 reciprocally reinforce their own transcription to establish an extensive circuit that regulates meiosis. TCFL5 promotes transcription of genes required for mRNA turnover, pachytene piRNA production, meiotic exit, and spermiogenesis. This transcriptional architecture is conserved in rhesus macaque, suggesting TCFL5 plays a central role in meiosis and spermiogenesis in placental mammals. *Tcfl5^em1/em1^* mutants are sterile, and spermatogenesis arrests at the mid- or late-pachytene stage of meiosis.

---

Gametogenesis converts diploid progenitor germ cells into haploid gametes specialized for sexual reproduction. Male gametogenesis encompasses a stepwise developmental pathway of meiotic cell divisions and differentiation to generate mature, swimming sperm from self-renewing progenitor germ cells^1^. Because haploid sperm are transcriptionally inert^2–4^, the meiotic cells must express not only gene products required for the orderly progression and completion of meiosis, but also produce the transcripts encoding proteins required for constructing mature sperm^1, 5^. Precise regulation of the meiotic gene expression program ensures male fertility^6–10^.

DNA replication in spermatogonia produces cells with 2N, 4C DNA content that differentiate into primary spermatocytes. These cells then undergo a lengthy meiotic I prophase, during which homologous chromatids pair and recombine to yield cells with 1N, 2C DNA content, the secondary spermatocytes. Meiosis II gives rise to haploid germ cells with 1N, 1C DNA content that differentiate into mature sperm, a process known as spermiogenesis (reviewed in ref. 1).

In human and mouse testes, ∼100 genes—first expressed at the pachytene stage of meiosis I—make non-coding transcripts that are processed into 26–30 nt pachytene PIWI-interacting RNAs (pachytene piRNAs)^11, 12^. Found only in placental mammals, pachytene piRNAs are diverse—comprising ∼1 million distinct species—and plentiful—rivaling ribosomal abundance^11–13^. Pachytene piRNAs bind the PIWI proteins MIWI and MILI, generating RNA-protein complexes that can repress transcripts during male gametogenesis^14–24^. Spermiogenesis requires pachytene piRNAs, yet their sequences are poorly conserved and rapidly diverging even among modern humans^12, 19, 24^.

The transcription factor (TF) A-MYB binds the promoters of more than half of pachytene piRNA-producing genes (“pachytene piRNA genes”) in humans and mice, and *A-Myb* mutant mice make fewer pachytene piRNAs than wild-type^11, 12^. Without A-MYB, germ cells fail to express meiotic genes and arrest early in meiosis I (ref. 5). Although A-MYB plays a crucial role regulating male meiosis, human and mouse testes with little or no A-MYB still make some pachytene piRNAs, and the promoters of half of human and macaque piRNA genes are not bound by A-MYB^12^. Moreover, A-MYB binds to the promoters of less than one-quarter of meiotic genes^5, 11^. These observations suggest that additional TFs are required to regulate male meiosis.

Here, we report that the testis-specific TF TCFL5 regulates meiosis. A-MYB initiates transcription of *Tcfl5* early in meiosis I. Subsequently, TCFL5 binds both its own promoter and that of *A-Myb*, establishing a mutually reinforcing positive feedback loop. Without TCFL5, male mouse germ cells arrest at the mid- or late-pachynema. Intriguingly, *Tcfl5* is haploinsufficient: *Tcfl5^+/em^*^1^ mice produce few sperm and are subfertile. Whereas A-MYB generally initiates the production of pachytene piRNAs from evolutionarily older piRNA precursor loci, transcription of younger pachytene piRNA genes is more dependent on TCFL5. Like A-MYB, TCFL5 promotes transcription of both piRNA precursors and mRNAs encoding piRNA biogenesis proteins. Consequently, the mutually reinforcing A-MYB/TCFL5 positive feedback loop allows each protein to initiate interlocking feed-forward loops dedicated to piRNA production. This transcriptional architecture is evolutionarily conserved in the old-world monkey rhesus macaque, suggesting it predates the divergence of rodents and primates. Our findings identify a transcriptional circuit that regulates meiotic gene expression and likely co-evolved with pachytene piRNAs themselves.

## Results

### TCFL5, a testis-specific TF first expressed at the pachytene stage of meiosis I

In mice, *Tcfl5* is specifically expressed in testis (Fig. 1a). We used staged mouse testes^25^ to measure the abundance of *Tcfl5* mRNA and protein across spermatogenesis. At 11 days post-partum (dpp), meiosis proceeds no further than the zygotene stage; early pachytene spermatocytes appear at 14 dpp; and mid- and late-pachytene spermatocytes are present by 17 dpp. *A-MYB* mRNA and protein were first detected at 14 dpp. By contrast, *Tcfl5* mRNA was present at 14 dpp, but TCFL5 protein was not detected until 17 dpp (Fig. 1b,c, and Extended Data Fig. 1a).

**Figure 1.**
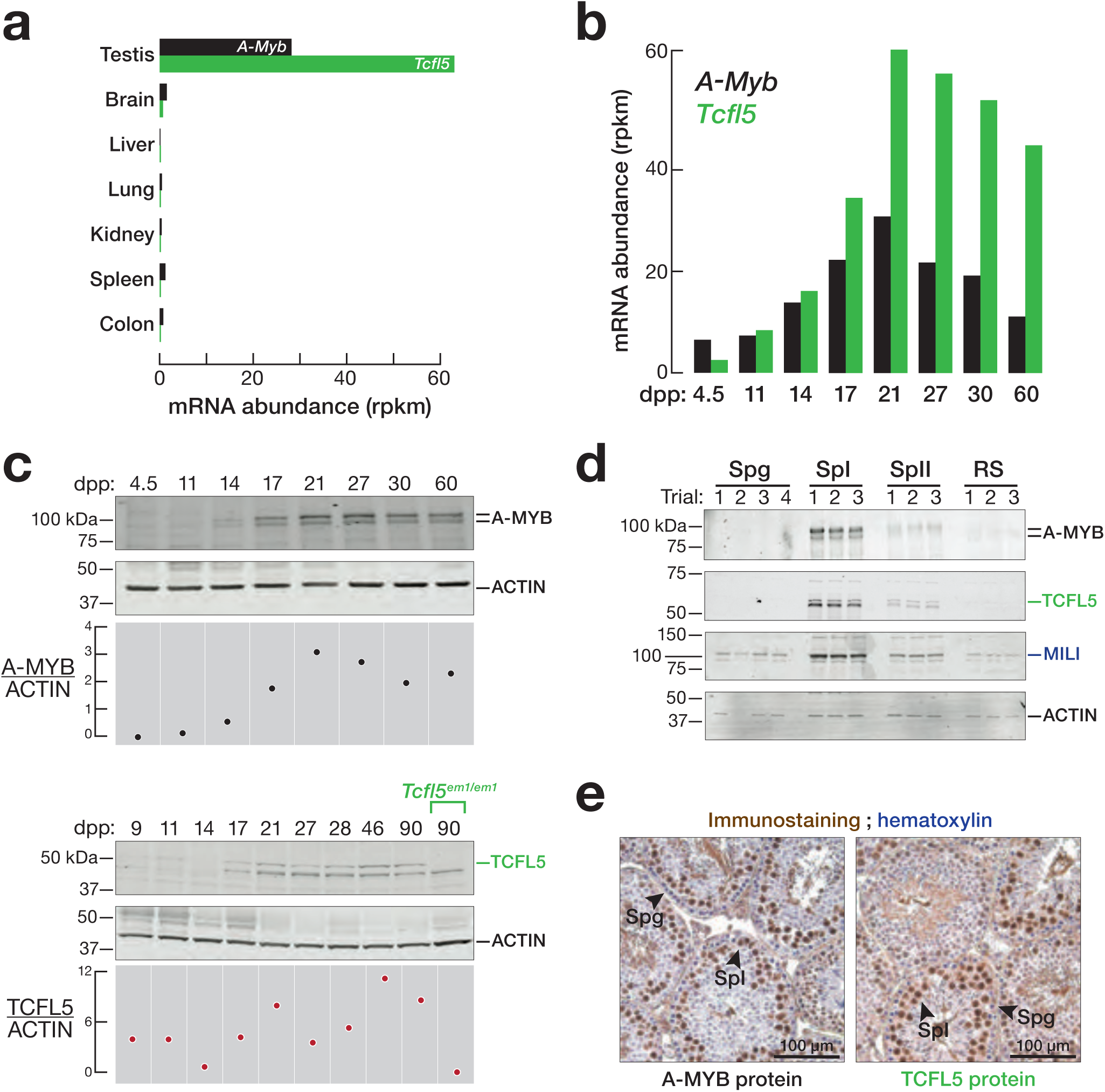
TCFL5 is expressed after A-MYB. **a,** Abundance of *A-Myb* and *Tcfl5* mRNA in various mouse tissues. **b,c,** *A-Myb* and *Tcfl5* mRNA (**b**) and protein abundance (**c**) at distinct developmental stages of mouse testis. Each lane contained 50 µg protein. **d,** Abundance of A-MYB and TCFL5 proteins in FACS-purified germ cells. Each lane contained protein from 100,000 germ cells. To detect TCFL5, the membrane was incubated with rabbit anti-TCFL5 (Sigma, HPA055223; 1:1,000). Spg: spermatogonia; SpI: primary spermatocytes; SpII: secondary spermatocytes; RS: round spermatids. **e,** Immunohistochemical detection of A-MYB and TCFL5 in adult mouse testis sections.

To directly test whether TCFL5 is expressed in primary spermatocytes, we measured TCFL5 protein in FACS-purified germ cells. MILI, which is present in both spermatogonia and spermatocytes, served as a control^11–14, 26–28^. A-MYB and TCFL5 were not detectable in purified spermatogonia, were abundant in primary spermatocytes (comprising pachytene and diplotene spermatocytes), and present at a lower level in secondary spermatocytes (Fig. 1d, Extended Data Fig. 1b). Immunostaining of A-MYB and TCFL5 in testis sections detected the two proteins in the nuclei of primary spermatocytes, but not in those of spermatogonia (Fig. 1e).

*A-Myb* expression precedes that of *Tcfl5*: dual-color RNA fluorescent in situ hybridization (RNA-FISH) on adult testis sections revealed that *A-Myb* mRNA first appears in the nuclei of leptotene/zygotene cells as one or two brightly staining loci, suggesting they correspond to the genomic sites of transcription (Fig 2a,b). *Tcfl5* mRNA was not detected in leptotene/zygotene cells, and first appeared in the nuclei and cytoplasm of pachytene cells (Fig 2a,b). Single-color RNA in situ hybridization of multiple seminiferous tubules at different stages confirmed that *A-Myb* mRNA first appears in pre-leptotene, leptotene, and zygotene cells, whereas *Tcfl5* first appears in mid- and late-pachytene cells (Extended Data Fig. 2a).

**Figure 2.**
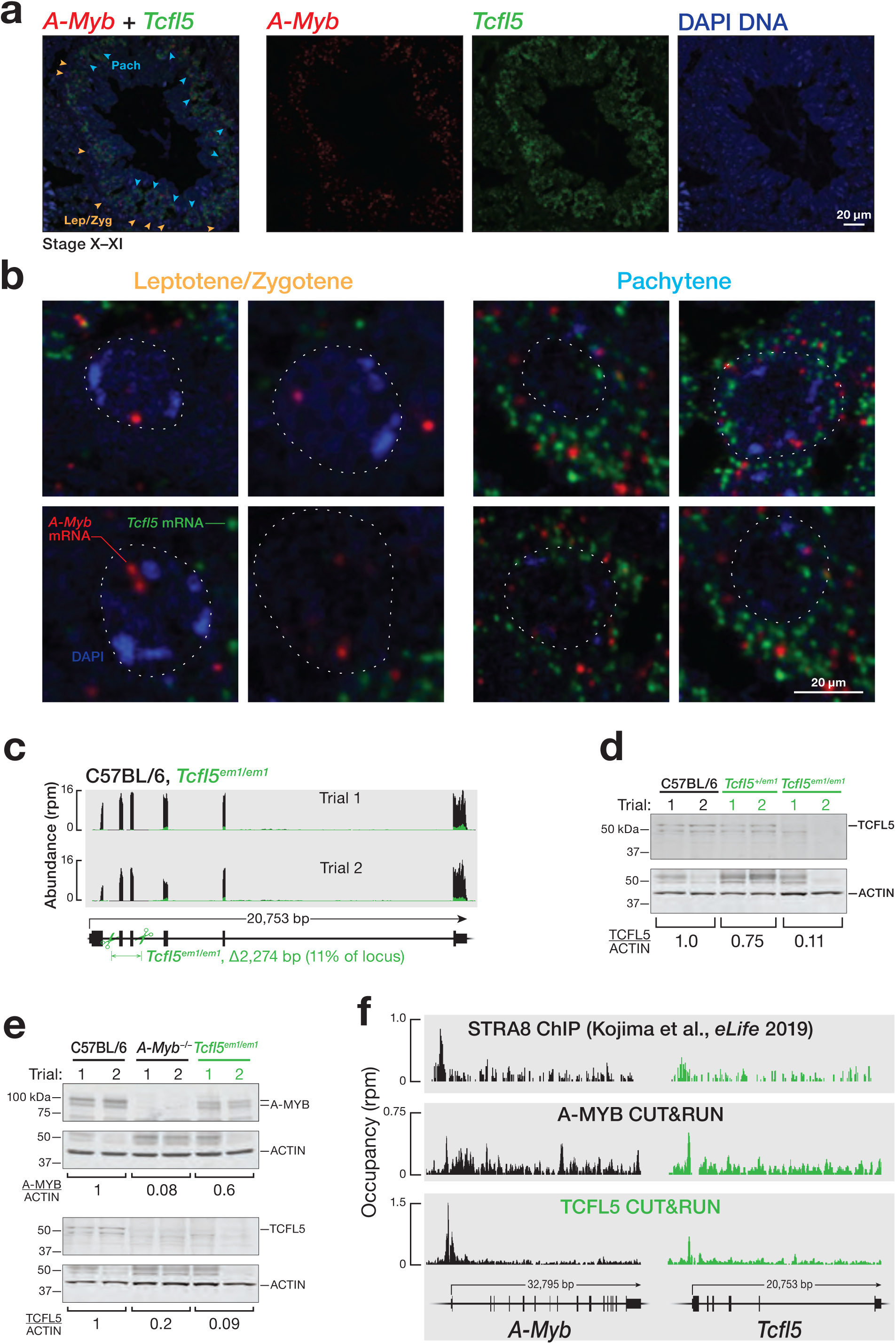
*A-Myb* expression precedes that of *Tcfl5*. **a,** mRNA expression of *A-Myb* and *Tcfl5* in a seminiferous tubule section from adult mouse testis, detected by two-color RNA fluorescent *in situ* hybridization. **b,** Representative images of individual cells from the experiment in (**a**), showing leptotene/zygotene- and pachytene-stage nuclei. **c,** Strategy to generate *Tcfl5* knockout mice using CRISPR/Cas9. Scissors indicate sites targeted by sgRNAs designed to delete *Tcfl5* exons 2 and 3. RNA-seq was used to measure the abundance of *Tcfl5* mRNA in adult testes. **d,** Protein abundance of TCFL5 in *Tcfl5^em1/em1^* mutant testes compared to C57BL/6 wild-type testes. ACTIN serves as a loading control. Each lane contained 50 µg testis lysate protein. **e,** Abundance of A-MYB and TCFL5 proteins in *A-Myb^−/−^* and *Tcfl5^em1/em1^* mutant mice testes. ACTIN serves as a loading control. Each lane contained 50 µg testis lysate protein. **f,** STRA8 ChIP-seq and A-MYB and TCFL5 CUT&RUN peaks at the promoters of *A-Myb* and *Tcfl5*.

### A-MYB initiates a reciprocal positive feedback loop between TCFL5 and A-MYB

*Tcfl5* mRNA was not detectable by RNA-FISH in *A-Myb^repro9/repro9^* (henceforth, *A-Myb^−/−^*) mutant testis (Extended Data Fig. 2b). We used a pair of single-guide RNAs (sgRNAs) to generate a *Tcfl5* knockout by deleting *Tcfl5* exons 2 and 3 (2,274 bp) (Fig. 2c,d and Extended Data 2c). *A-Myb* mRNA was readily detected in *Tcfl5^em1Pdz/em1Pdz^* (henceforth, *Tcfl5^em1/em1^*) (Extended Data Fig. 2b). A-MYB protein was present in *Tcfl5^em1/em1^*, but was less abundant than in C57BL/6 controls (Fig. 2e, Extended Data Fig. 2d). By contrast, the abundance of TCFL5 protein in *A-Myb^−/−^* mutant testes was indistinguishable from that in *Tcfl5^em1/em1^* mutants (Fig. 2e and Extended Data Fig. 2d). These data suggest that A-MYB initiates *Tcfl5* expression, while TCFL5 acts to increase the steady-state abundance of A-MYB.

To determine whether A-MYB regulates *Tcfl5* directly or indirectly, we performed cleavage under targets and release using nuclease (CUT&RUN^29^) on FACS-purified primary spermatocytes. A prominent A-MYB peak was located at the first nucleotide of the *Tcfl5* transcription start site (TSS) (Fig. 2f). TCFL5 CUT&RUN of FACS-purified primary spermatocytes revealed a prominent TCFL5 peak within the promoters of both *A-Myb* and *Tcfl5* itself (Fig. 2f). ChIP-seq of A-MYB and TCFL5 from adult mouse testes confirmed these observations (Extended Data Fig. 2e). We conclude that A-MYB promotes its own transcription and directly initiates transcription of *Tcfl5*; TCFL5 responds by reinforcing its own transcription and that of A-MYB via positive transcriptional feedback.

What turns on *A-Myb* expression? The retinoic acid-responsive TF STRA8 triggers the entrance of spermatogonia into meiosis I (ref. 30). Our re-analysis of publicly available STRA8 ChIP-seq data^30^ identified a prominent STRA8 peak on the promoter of *A-Myb*, but not *Tcfl5* (Fig. 2f). Thus, STRA8 initiates A-MYB expression upon entry into meiosis I, and that A-MYB subsequently drives transcription of *Tcfl5*.

### TCFL5-deficient germ cells arrest later than *A-Myb* mutants

*Tcfl5^em1/em1^* mutant testes lack epididymal sperm (Fig. 3a,b). Consistent with the testis-specific expression of *Tcfl5*, we observed no abnormalities in *Tcfl5^em1/em1^* mutant females. Gross histology of *Tcfl5^em1/em1^* mutant testis revealed that mutants arrest spermatogenesis at meiosis I and contain no spermatids or spermatozoa (Fig. 3b). Strikingly, the developing pachytene spermatocytes in *Tcfl5^em1/em1^* mutants formed abnormal multinucleated symplasts (Fig. 3b). These giant cells contain multiple nuclei within a common cytoplasm, suggesting incomplete cytokinesis. TUNEL staining showed that the majority of these giant multinucleated symplastic germ cells did not undergo apoptosis (Extended Data Fig. 3a). Immunostaining to detect the synaptonemal complex protein SYCP3 suggested that the chromosomes of these multinucleated pachytene spermatocytes were of normal length and appearance (Extended Data Fig. 3b,c). Consistent with TUNEL staining, immunostaining for γH2AX, a marker for DNA double-strand breaks (DSBs), detected only the normal enrichment of γH2AX on the transcriptionally inactive, asynapsed sex chromosomes^31^; no increase in autosomal DSBs as marked by γH2AX was observed (Extended Data Fig. 3b,c). *A-Myb^−/−^* pachytene spermatocytes feature cloud-like γH2AX staining over the autosomal chromosomes, consistent with unrepaired DSBs persisting in unsynapsed chromatin^5^; these abnormalities were not present in *Tcfl5^em1/em1^* mutants (Extended Data Fig. 3c).

**Figure 3.**
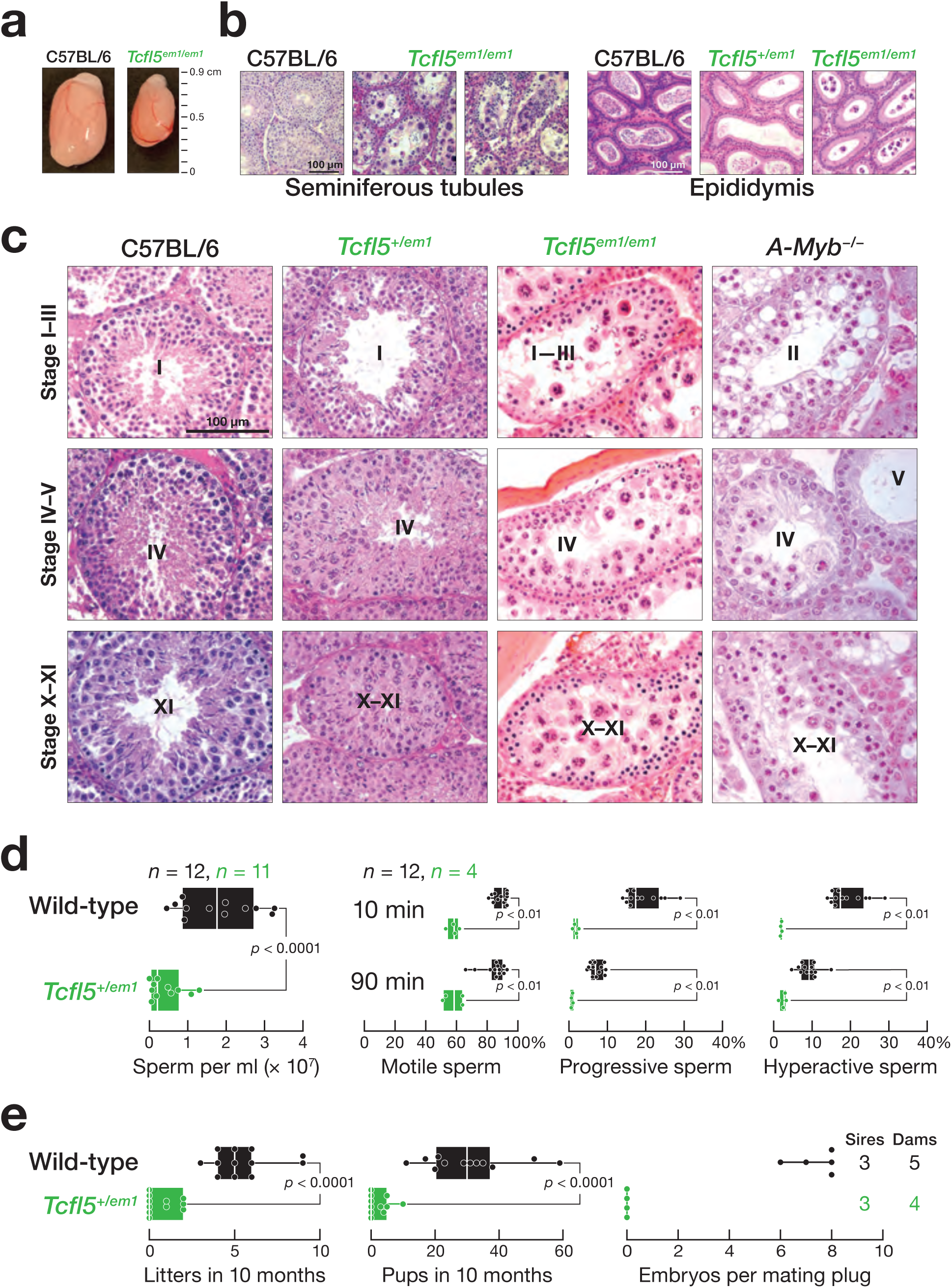
*Tcfl5^em1/em1^* mutant germ cells arrest at the mid-pachytene stage of meiosis I, and *Tcfl5^+/em1^* mutant mice fail to produce functional sperm. **a,** Comparison of testes from four-month-old C57BL/6 and *Tcfl5^em1/em1^* mice. **b, c,** Hematoxylin and eosin staining of (**b**) testis and epididymis sections from C57BL/6, *Tcfl5^em1/em1^*, and *Tcfl5^+/em1^* mice and (**c**) seminiferous tubules at different epithelial stages from C57BL/6, *Tcfl5^em1/em1^*, *Tcfl5^+/em1^*, and *A-Myb^−/−^* mice. **d,** Concentration and the percent motile, progressive and hyperactivated sperm isolated from *Tcfl5^em1/em1^* and C57BL/6 control caudal epididymis. **e,** Viable litters and total pups sired in 10 months by two-to-eight-month-old male mice, and the number of embryos present at 14.5 days post-coitus in C57BL/6 females mated to *Tcfl5^+/em1^* or C57BL/6 wild-type males. For (**d**) and (**e**), vertical lines denote median, and whiskers mark the minimum and maximum values. Each dot represents an individual male. Significance was measured using a two-tailed unpaired Mann-Whitney U test

Thus, A-MYB-deficient germ cells appear unable to complete synapsis, whereas *Tcfl5^em1/em1^* mutant germ cells complete synapsis and continue to progress through the pachynema, ultimately forming multinucleated symplasts that are unable to complete meiosis. Supporting this view, *A-Myb^−/−^* mutant testis have a loose, defective epithelium by stage II, whereas *Tcfl5^em1/em1^* mutants first show developmental defects at stage IV (Fig 3c). Stage-IV tubules from *A-Myb^−/−^* mutants contain few early pachytene cells; by stage V, *A-Myb^−/−^* mutant testis have lost almost all pachytene spermatocytes (Fig. 3c). By contrast, stage-IX tubules from *Tcfl5^em1/em1^* mutants contain late pachytene spermatocytes (Fig. 3c).

### TCFL5 is haploinsufficient

Testes from *Tcfl5^+/em1^* heterozygotes were not normal. *Tcfl5^+/em1^* testis sections showed fewer epididymal sperm than wild-type mice (Fig. 3b), and the quantity of caudal epididymal sperm produced by *Tcfl5^+/em1^* mice was ∼5-fold less than C57BL/6 mice (two-sided unpaired Mann-Whitney-Wilcoxon U test, *p* < 0.0001) (Fig. 3d). Compared to C57BL/6, *Tcfl5^+/em1^* mice had significantly fewer motile and progressive sperm, and very few sperm were hyperactivated when incubated in capacitating conditions (Fig. 3d). When mated with C57BL/6 females, adult *Tcfl5^+/em1^* males sired fewer viable litters and fewer pups than C57BL/6 males (two-sided unpaired Mann-Whitney-Wilcoxon U test, both *p* < 0.0001) (Fig. 3e). Moreover, the embryos sired by *Tcfl5^+/em1^* males often failed to develop. For example, three *Tcfl5^+/em1^* males were each housed with two C57BL/6 females, and embryos isolated 14.5 days after the appearance of a mating plug. No embryos were detected among the C57BL/6 females paired with *Tcfl5^+/em1^* males (*n* = 4 dams and 3 sires); females paired with C57BL/6 males all carried healthy embryos (7.4 ± 0.9, *n* = 5 dams and 3 sires) (Fig. 3e).

To better understand the underlying molecular defects that lead to male infertility in *Tcfl5^+/em1^* heterozygotes, we measured steady-state RNA abundance in primary spermatocytes purified from adult testes (Fig. 4a). The mRNA abundance of 4,653 genes in primary spermatocytes were significantly lower in *Tcfl5^+/em1^* compared with C57BL/6 (mutant ÷ control < 0.5 and FDR < 0.05) (Fig. 4a and Supplementary Table 1). Notably, the abundance of *Tcfl5* mRNA itself was fivefold lower in *Tcfl5^+/em1^* primary spermatocytes than in wild-type (mean change = 0.17 ± 0.06).

**Figure 4.**
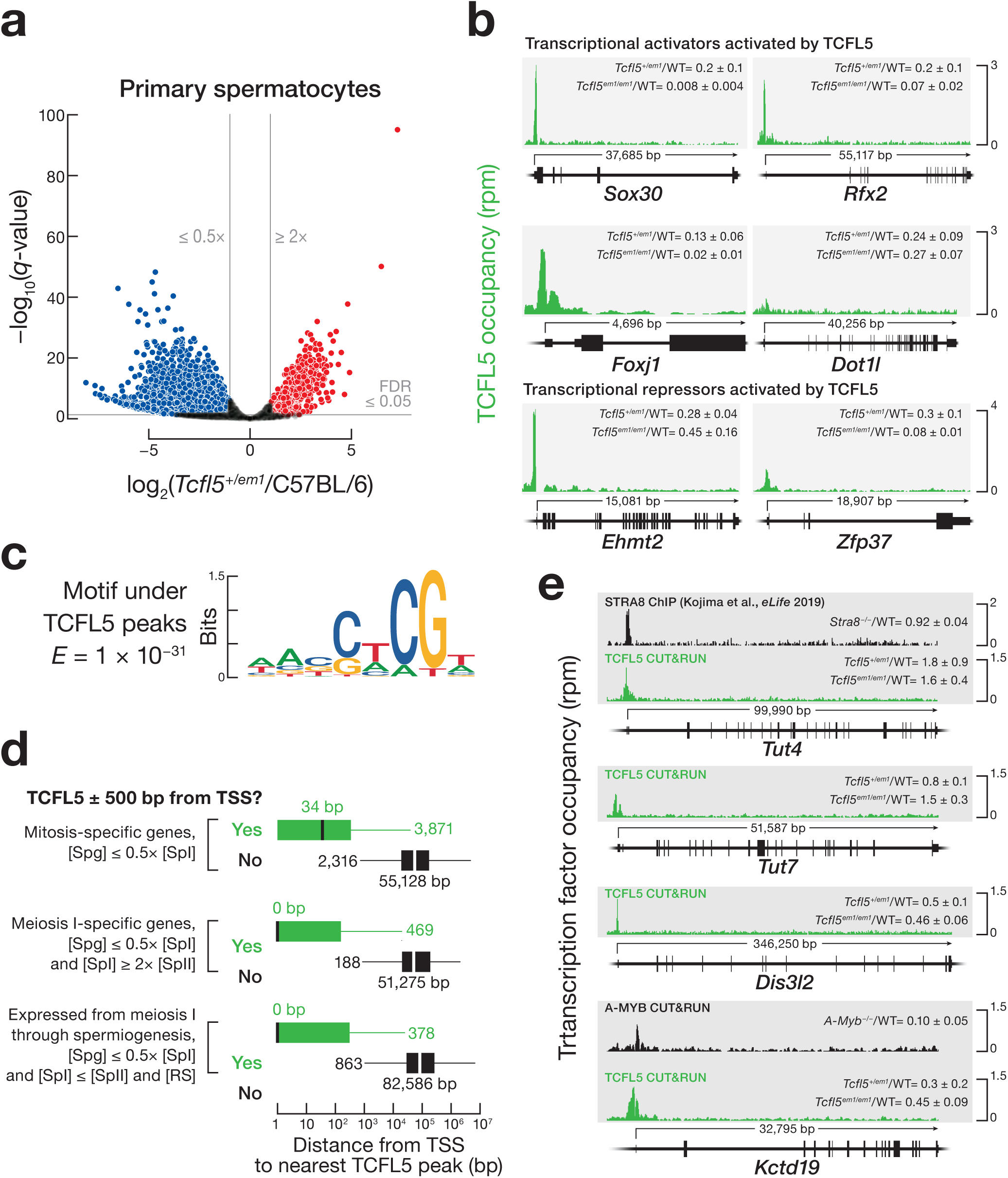
TCFL5 regulates male meiosis I and spermiogenesis. **a,** Volcano plot shows the significantly deregulated genes in FACS-purified primary spermatocytes from *Tcfl5^+/em1^* heterozygous testes (*n* = 3), compared to C57BL/6 (*n* = 3). **b,** TCFL5 CUT&RUN peaks at the promoters of *Sox30*, *Rfx2*, *Foxj1*, *Dot1l*, *Ehmt2*, and *Zfp37*. **c,** MEME-reported canonical TCFL5 binding motif beneath all CUT&RUN TCFL5 peaks. **d,** The mean distance of two replicates from the nearest TCFL5 peak determined by CUT&RUN to the transcription start site (TSS) for mitosis-specific genes, meiosis I-specific genes, and genes whose expression starts at meiosis and persists through spermiogenesis. Genes classified as not bound by TCFL5 showed no TCFL5 peak ±500 bp from the TSS in any of the CUT&RUN or ChIP-seq datasets. Vertical black lines: median; whiskers: maximum and minimum values, excluding outliers. **e,** STRA8 ChIP-seq and TCFL5 or A-MYB CUT&RUN peaks at the promoters of *Tut4/7*, *Dis3l2*, and *Kctd19*. In (**b**), and (**e**), WT denotes C57BL/6.

Many of these changes likely reflect a direct role of TCFL5 in the transcriptional activation of genes whose mRNA abundance was reduced in *Tcfl5^+/em1^*. CUT&RUN analysis of TCFL5 DNA binding revealed that TCFL5 bound near the transcription start site (TSS) for 2,173 of 4,653 genes whose transcripts were reduced in *Tcfl5^+/em1^* primary spermatocytes (Extended Data Fig. 4a).

*Tcfl5* haploinsufficiency made obtaining *Tcfl5^em1/em1^* males daunting: two homozygous males survived to sexual maturity among eight breeding pairs over 2.5 years. Moreover, adult *Tcfl5^em1/em1^* whole testes contain a higher fraction of primary spermatocytes than wild-type, so comparisons of transcript expression between *Tcfl5^em1/em1^* and wild-type testes underestimate decreases in TCFL5-dependent gene expression. Therefore, our conclusions are based primarily on comparison of gene expression in purified primary spermatocytes from *Tcfl5^+1/em1^* heterozygous mutants and C57BL/6 controls.

### TCFL5 activates transcription of transcription factor genes

Of the 1,304 genes whose steady-state transcript abundance was reduced more than twofold in *Tcfl5^em1/em1^* mutant mice whole testes compared to wild-type (median change = 0.2 ± 0.1) and whose promoters were bound by TCFL5, the transcript abundance of 1,058 was also reduced in *Tcfl5^+/em1^* primary spermatocytes (median change = 0.26 ± 0.09). Of these 1,058 genes, the promoters of only ∼42% (443 of 1,058) were occupied by A-MYB, suggesting that >600 genes are transcriptionally activated by TCFL5 rather than A-MYB.

Beyond those genes directly activated by TCFL5, many others are likely regulated by TFs that are themselves under TCFL5 control. In fact, 73 chromatin modifying enzymes or sequence-specific, DNA-binding transcription factors whose promoters are bound by TCFL5, showed significantly reduced mRNA abundance in both *Tcfl5^+/em1^* primary spermatocytes and *Tcfl5^em1/em1^* mutant testes compared to wild-type, strong evidence that they require TCFL5 for their activation (Extended Data Fig. 4b and Supplementary Table 1). These include *Sox30*, *Rfx2*, and *Foxj1*, which encode transcription factors indispensable for spermiogenesis, as well as *Dot1l*, which encodes a conserved methyltransferase that methylates lysine 79 in histone H3 at a subset of enhancer elements and regulates homologous recombination during meiosis in *C. elegans*^32–34^ (Fig. 4b). SOX30 is required for male fertility, and both SOX30 and RFX2 activate the expression of genes that participate in spermiogenesis^35–37^. The forkhead family TF FOXJ1 activates the transcription of ciliary genes^38, 39^ and regulates sperm motility in frog testis^40^. In addition to these transcriptional activators, the 73 TCFL5-regulated transcription factor genes include two repressors, EHMT2 and ZFP37 (refs. 41,42) (Fig. 4b). Dimethylation of lysine 9 of histone H3 by EHMT2 has been proposed to ensure that recombination occurs at the correct genomic loci^41^.

### Mapping TCFL5 binding across the genome

We identified genes with high TCFL5 binding occupancy using Sparse Enrichment Analysis for CUT&RUN (SEARC^43^) data from FACS-purified primary spermatocytes. SEARC identified 9,290 genes, comprising 7,664 protein-coding and 332 non-coding genes, with a TCFL5 peak within 500 bp of their transcription start site (TSS) (Supplementary Table 3). The sequence motif associated with the CUT&RUN peaks defined the TCFL5 consensus binding site as WANSWCGW (W = A or T; S = G or C; expected value by chance, *E* = 1 × 10^−31^) (Fig. 4c). ChIP-seq of TCFL5 using whole adult testis corroborated the CUT&RUN data from sorted cells (Extended Data Fig. 4d and Supplementary Table 3).

To better understand the role of TCFL5 in regulating meiotic gene expression, we classified genes according to their transcript concentration: i.e., molecules per cell corrected for cell volume (Extended Data Fig. 4c and Supplementary Table 2a,b). Mitosis-specific genes were defined as those whose transcript concentration in spermatogonia was more than twice that in primary spermatocytes (8,209 genes). Meiosis I-specific genes were required to have a transcript concentration in primary spermatocytes more than twice that in either spermatogonia or secondary spermatocytes (792 genes). mRNAs required during spermiogenesis are mostly transcribed during meiosis^2–4^. A third category comprised genes turned on at meiosis I and whose mRNAs persisted through spermiogenesis; these were defined as having a transcript concentration in primary spermatocytes that was more than twice that in spermatogonia but was essentially unchanged in secondary spermatocytes and round spermatids (1,466 genes).

The promoters of ∼47% of mitosis-specific genes were bound by TCFL5 (3,871 of 8,209 genes; median distance from TSS to the nearest TCFL5 peak = 34 bp), suggesting TCFL5 sustains their expression after the onset of meiosis I (Fig. 4d and Supplementary Table 3). Approximately 59% of the promoters of the meiosis I-specific genes were bound by TCFL5 (469 of 792 genes; median distance from TSS to the nearest TCFL5 peak = 0 bp); these genes are likely turned on or up by TCFL5 early in meiosis I Among the 1,466 genes expressed at meiosis I whose expression persisted through spermiogenesis, TCFL5 bound to the promoters of ∼26% (378 of 1,466 genes; median distance from TSS to the nearest TCFL5 peak = 0 bp).

Notably, TCFL5 binds the promoters of *Tut4* (*Zcchc11*), *Tut7* (*Zcchc6*), and *Dis3l2* (Fig. 4e). This pair of terminal nucleotide transferases uridylates zygotene-expressed mRNAs whose 3′ untranslated regions contain AU-rich elements (AUUUA; AREs), marking them for destruction by the 3′-to-5′ exoribonuclease DIS3L2 as cells transit from the zygotene to the pachytene stage^44^. The mRNA abundance of *Tut7* and *Dis3l2* increase >5-fold in primary spermatocytes, compared to spermatogonia, whereas *Tut4* increases only modestly (∼1.6-fold). The expression pattern of *Tut7* and *Dis3l2* suggests that these genes are activated by the TCFL5 bound to their promoters. Yet, the mRNA abundance of *Tut4* increased ∼50% in both *Tcfl5^em1/em1^* mutant mice and *Tcfl5^+/em1^* primary spermatocytes, whereas *Dis3l2* mRNA decreased twofold (*Tut4*: mean increase = 1.6 ± 0.4 in *Tcfl5^em1/em1^* whole testes and 1.8 ± 0.9 in *Tcfl5^+/em1^* primary spermatocytes; *Dis3l2*: mean decrease = 0.46 ± 0.06 in *Tcfl5^em1/em1^* whole testes and 0.5 ± 0.1 in *Tcfl5^+/em1^* primary spermatocytes, relative to C57BL/6) (Fig. 4e). Although we cannot exclude the possibility that TCFL5 represses expression of *Tut4* but activates *Dis3l2*, we propose an alternative explanation. TCFL5 likely activates the expression of all three, but the expected decrease of *Tut4* and *Tut7* mRNA in *Tcfl5* mutants is obscured by an increase in their mRNA stability because *Tut4* and *Tut7* mRNAs are themselves ARE-dependent targets of TUT4/TUT7-catalyzed uridylation and DIS3L2 degradation. Consequently, in the absence of TCFL5, when *Dis3l2* expression decreases, *Tut4* and *Tut7* mRNAs become more stable. Supporting this view, the *Tut4* 3′ UTR contains three canonical AREs, and the 1030-nt long *Tut7* 3′ UTR both contains one canonical ARE and is longer than typical for testis (median = 639 nt), a feature shared by mRNAs targeted by TUT4/7 targets (median = 1,703 nt) generally^44^.

Finally, the set of promoters bound by TCFL5 includes *Kctd19* (Fig. 4e), a gene whose protein product is essential for meiotic exit^45^. The *Kctd19* promoter is also bound by A-MYB. The abundance of *Kctd19* mRNA increases >200-fold when spermatogonia differentiate into spermatocytes (Supplementary Table 2a). *Kctd19* mRNA abundance declines >3-fold in *Tcfl5^+1/em1^* and 10-fold in *A-Myb^−/−^* primary spermatocytes.

### TCFL5 regulates the expression of piRNA pathway genes

At the onset of the pachynema in mice and humans, A-MYB activates transcription of genes producing pachytene piRNA precursors and genes encoding piRNA biogenesis proteins, thereby generating a feedforward loop that drives pachytene piRNA production^11, 12^. In parallel, TCFL5 also establishes a feedforward loop that powers pachytene piRNA production.

In FACS-purified primary spermatocytes, the promoters of 14 piRNA biogenesis genes featured a TCFL5 peak near their TSSs (median distance = 0 bp) (Fig. 5a and Extended Data Fig. 5a). The steady-state mRNA abundance of five of the TCFL5-bound piRNA biogenesis genes—*Gtsf1*, *Henmt1*, *Mael, Pld6*, and *Tdrd6—*was reduced ≥2-fold in *Tcfl5^em1/em1^* and *Tcfl5^+1/em1^* testes (Extended Data Fig. 5a,b). Among the remaining nine piRNA biogenesis genes, the mRNA abundance of five decreased modestly in *Tcfl5^+1/em1^*. The promoters of the nine genes were all bound by A-MYB, suggesting that their transcription is either driven entirely by A-MYB or that A-MYB largely compensates in *Tcfl5* mutant mice.

**Figure 5.**
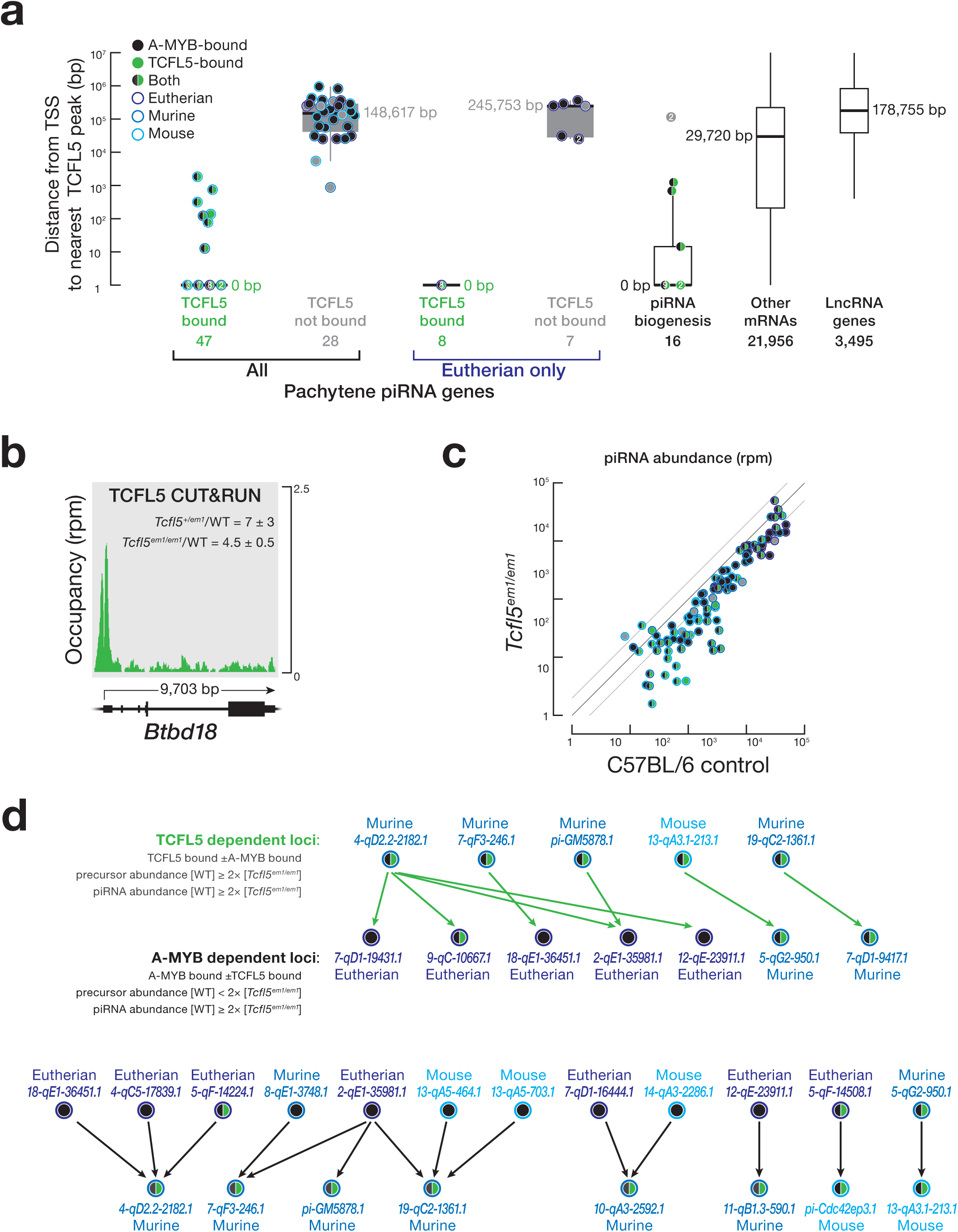
Younger pachytene piRNA genes activated by TCFL5 produce piRNAs that initiate piRNA production in older pachytene piRNA genes. **a,** The distance from the nearest TCFL5 peak to the transcription start sites (TSS) of pachytene piRNA genes, genes encoding piRNA biogenesis proteins, and other protein-coding or non-coding genes. Genes were classified as not bound by TCFL5 as in Fig. 4. Horizontal lines: median; whiskers: maximum and minimum values, excluding outliers. Each dot represents mean distance of two replicates of CUT&RUN from the nearest TCFL5 peak to the TSS of individual genes. Measurements with the same values are indicated by a single marker indicating number of individual data points. *Gpat2*, which is required for piRNA biogenesis in flies^64^ and in cultured mouse germline stem cells^65^, was not included in this analysis, because its participation in the piRNA pathway in vivo in mice remains to be established. **b,** TCFL5 CUT&RUN peak at the promoter of *Btbd18*. WT denotes C57BL/6. **c,** Steady-state abundance of pachytene piRNAs in C57BL/6 and *Tcfl5^em1/em1^* mice. Each dot represents a pachytene piRNA gene. **d,** Pachytene piRNA-directed cleavage sites in pachytene piRNA precursor transcripts. *Top*: each arrow points from the piRNAs from five pachytene piRNA genes whose expression was reduced >2-fold in *Tcfl5^em1/em1^* mutant mice towards the inferred cleavage sites in seven pachytene piRNA precursor transcripts whose transcription is activated by A-MYB. *Bottom*: each arrow points from the piRNAs from 12 pachytene piRNA genes whose expression was essentially unchanged in *Tcfl5^em1/em1^* mutant mice, towards eight pachytene piRNA precursor transcripts whose abundance was reduced >2-fold in *Tcfl5^em1/em1^* mutant mice. Transcription of these 12 pachytene piRNA genes is activated by A-MYB.

### TCFL5 activates evolutionarily younger pachytene piRNA genes

Our data suggest that TCFL5 activates transcription of almost half of pachytene piRNA genes during male meiosis I. CUT&RUN of FACS-purified primary spermatocytes identified TCFL5 peaks within 500 bp of the TSSs of 47 of the 100 mouse pachytene piRNA genes (median distance = 0 bp), whereas 91 were bound by A-MYB (median distance = 0 bp0 (Fig. 5a and Supplementary Table 4).

Pachytene piRNA genes with evolutionary conservation of genomic location (synteny) have first exons longer than 10 kbp and a low level of CG dinucleotides; their promoters and gene bodies are bound by the transcription elongation factor BTBD18 and marked by histone lysine acylation, including acetylation, butyrylation, and crotonylation^46, 47^. BTBD18 encodes a transcriptional elongation factor that has been proposed to facilitate the production of piRNA precursors from evolutionarily older pachytene piRNA genes^46^, and precursor transcript and mature piRNA abundance from these loci declines significantly in *Btbd18* mutant testes^47^. *Btdbd18* mRNA abundance is highest in spermatogonia, and declines >12-fold in primary spermatocytes (Supplementary Table 2a), suggesting the protein marks pachytene piRNA loci prior to meiosis I. TCFL5 binds the *Btbd18* promoter (distance from TSS to the nearest TCFL5 peak = 0 bp), and *Btbd18* mRNA increased in both *Tcfl5^em1/em1^* mutant testes (mean increase = 4.5 ± 0.5) and *Tcfl5^+/em1^* primary spermatocytes (mean increase = 7 ± 3) compared to wild-type (Fig. 5b). We do not yet know whether TCFL5 serves to repress BTBD18 expression. Increased BTBD18 levels in mutants may help explain why loss of TCFL5 has little effect on the transcription of many evolutionary older pachytene piRNA genes.

A-MYB occupancy was greatest at the promoters of pachytene piRNA genes with low CG dinucleotide content (Spearman’s *ρ* = *−*0.35; Extended Data Fig. 6a), long first exons (*ρ* = 0.28), high histone lysine acylation (*ρ* = 0.38–0.63) and BTBD18 occupancy (*ρ* = 0.44; Extended Data Fig. 6a). High A-MYB and low TCFL5 promoter occupancy was also associated with pachytene piRNA genes producing abundant piRNAs (Extended Data Fig. 6b). By contrast, TCFL5 occupancy was correlated with high CG dinucleotide promoter sequence (Spearman’s *ρ* = 0.67; Extended Data Fig. 6a), a feature more likely to be found in evolutionarily younger pachytene piRNA genes, which typically produce fewer piRNAs^46^.

Strikingly, TCFL5 binds the promoters of seven genes encoding histone acyl transferases (reviewed in ref. 48). Of these seven genes, four were also bound by A-MYB (Extended Data Fig. 6c). Histone acyl transferases use acetyl-, butyryl, or crotonyl-CoA as acyl donors. Histone crotonylation, which promotes gene expression, is regulated by the intracellular concentration of crotonyl-CoA^49^. *Acss2* encodes a short chain acyl-CoA synthetase^50^ that uses ATP to synthesize acetyl- and crotonyl-CoA. TCFL5 binds the *Acss2* promoter.

Our data suggest that pachytene piRNA genes with high A-MYB occupancy are more likely to have conserved synteny than those with high TCFL5 and low A-MYB occupancy (Supplementary Table 4). Mouse pachytene piRNA genes were generally present at syntenic sites in other placental mammals (eutheria) when their promoters had high A-MYB occupancy (median = 3.8 RPM, 95% confidence interval [CI] = 2.8–4.7) compared to loci found only in murine rodents (median = 2.6 RPM, 95% CI = 2.1–3.2) or only in mouse (median = 2.3 RPM, 95% CI = 1.6–3.2) (Extended Data Fig. 6d and Supplementary Table 4). Conversely, TCFL5 promoter occupancy was higher for murine (median = 1.0 RPM, 95% CI = 0.6–2.3) and mouse pachytene piRNA genes (median = 1.3 RPM, 95% CI = 0.4–2.1) than at eutherian loci (median = 0.8 RPM, 95% CI = 0.5–1.1). Finally, the promoters of 20 of the 21 eutherian pachytene piRNA genes were bound by A-MYB, but only eight were occupied by TCFL5 (Fig. 5a). Our finding that A-MYB activates transcription of older pachytene piRNA genes, whereas TCFL5 tends to activate younger loci, suggests that acquisition of a TCFL5-binding site may facilitate the evolution of new pachytene piRNA genes.

### *Tcfl5* mutants have fewer pachytene piRNAs

For the majority of pachytene piRNA genes (77 of 100), piRNA abundance in *Tcfl5^em1/em1^* testes was less than half that in C57BL/6 controls (Fig 5c).The affected pachytene piRNA genes include loci with either or both A-MYB or TCFL5 bound to their promoters, as well as five bound by neither TF. The loss of pachytene piRNAs in *Tcfl5* mutants likely reflects the combined effects of reduced transcriptional activation by TCFL5 and A-MYB and reduced expression of pachytene piRNA biogenesis proteins. Loss of initiator piRNAs may also contribute to reduced piRNA levels.

piRNA production begins when a PIWI protein, loaded with an initiator piRNA, cleaves a complementary piRNA precursor to generate a 5′ monophosphorylated pre-pre-piRNA. Binding of a second PIWI protein to the 5′ end of the pre-pre-piRNA allows it to be further processed into a PIWI-loaded responder piRNA that can initiate additional piRNA production at fully or partially complementary sites^13, 18, 22–24, 51, 52^. Do piRNAs from TCFL5-activated pachytene piRNA genes initiate production of piRNAs from precursors whose transcription depends solely on A-MYB?

To test the idea that production of A-MYB-dependent piRNAs requires initiation by piRNAs from loci whose transcription is activated by TCFL5, we identified A-MYB-dependent piRNA precursor transcripts whose precursor transcript abundance declined less than twofold in in *Tcfl5^em1/em1^* but whose piRNA abundance declined twofold or more, suggesting a defect in piRNA biogenesis rather than transcriptional activation. Among these, we search for piRNAs from TCFL5-dependent pachytene piRNA genes capable of directing cleavage of the A-MYB-dependent precursor transcripts. Candidate piRNAs were required (1) to have complete seed complementarity (g2–g7) to the A-MYB-dependent precursor RNA; (2) form at least ten additional base pairs within the region g8–g21 with that precursor; and (3) be able to generate the 5′ end of a piRNA by PIWI-protein-catalyzed precursor RNA cleavage. The 5′ ends of 29 piRNAs from five different TCFL5-dependent pachytene piRNA precursors are predicted to target seven A-MYB-dependent pachytene piRNA precursors. Four of the five TCFL5-dependent pachytene piRNA genes are found among murine rodents and one was found only in mouse; none are conserved among eutheria generally. By contrast, five of the seven pachytene piRNA genes targeted by piRNAs from TCFL5-dependent pachytene piRNA genes are present at syntenic sites in other eutherian species; the remaining two are found in other murine rodents (Fig. 5d and Supplementary Table 5). Conversely, 62 piRNAs from 12 A-MYB-dependent pachytene piRNA genes are predicted to target the precursors from eight TCFL5-dependent pachytene genes. Seven of the 12 A-MYB-dependent pachytene piRNA genes are present at syntenic sites in other eutheria, two are found among murine rodents, while three were found only in mouse. Among the eight TCFL5-dependent pachytene piRNA genes targeted by these 12 loci, none were present in non-murine species (Fig. 5d and Supplementary Table 5).

### The regulatory architecture that drives pachytene piRNA production in rodents is conserved in rhesus macaque

In both rodents and primates, A-MYB activation of piRNA pathway proteins and pachytene piRNA genes creates a coherent feedforward loop that ensures the robust accumulation of pachytene piRNAs^11, 12^. In mouse, TCFL5 similarly organizes a coherent feedforward loop that activates the production of pachytene piRNAs.

In addition to placental mammals, TCFL5 orthologs are found in the genomes of marsupials, monotremes, birds, reptiles, amphibians, and fishes (Fig. 6a). Thus, TCFL5 likely participates in spermatogenesis across vertebrate species. Moreover, the A-MYB/TCFL5 regulatory architecture—reciprocal positive feedback circuits that reinforce *A-Myb* and *Tcfl5* transcription, sitting atop parallel A-MYB and TCFL5 coherent feedforward loops that drive pachytene piRNA production—is conserved in rhesus macaque (Fig. 6b,c and Supplementary Table 6). ChIP-seq of TCFL5 from adult rhesus testis identified 17,959 genomic regions with significant TCFL5 peaks (FDR < 0.05), comprising 4,048 annotated genes. The TCFL5-bound genes include six of the 15 rhesus orthologs of known piRNA biogenesis proteins and 30 of the 189 rhesus piRNA genes^12^ (median distance from TSS to nearest TCFL5 peak = 0 bp) (Fig 6c). Together, these data suggest that the A-MYB/TCFL5 transcriptional architecture dates to the last common ancestor of placental mammals.

**Figure 6.**
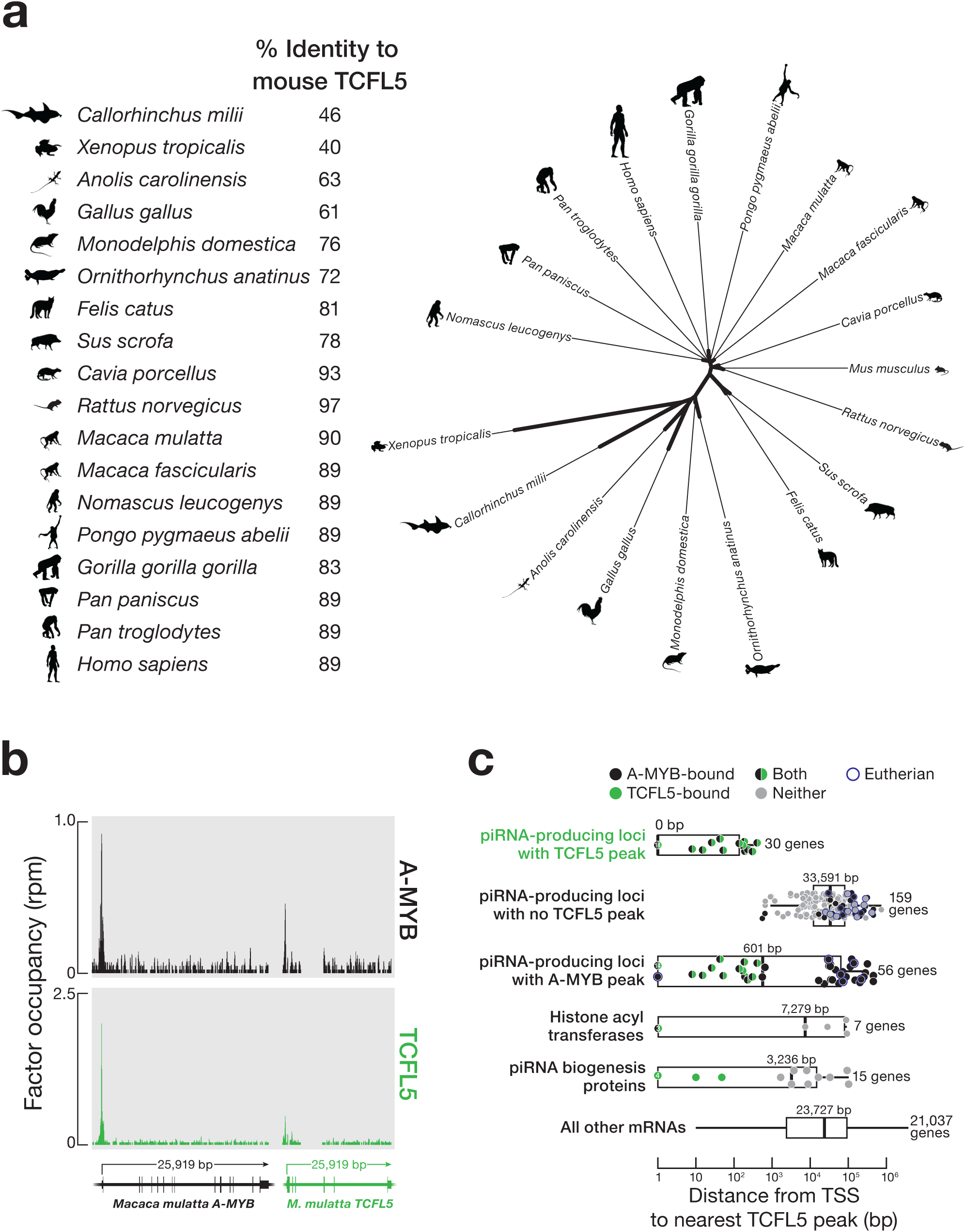
piRNA production by TCFL5 is conserved in rhesus macaque. **a,** Left: Percent identity of mouse TCFL5 to TCFL5 in other mammals. Right: Unrooted tree of TCFL5 protein sequence, aligned using Clustal Omega; unrooted tree was constructed using Randomized Axelerated Maximum Likelihood with default parameters and visualized in Interactive Tree Of Life. **b,** A-MYB and TCFL5 ChIP-seq peaks at the promoters of the rhesus macaque *A-MYB* and *TCFL5* genes. **c,** The distance from the nearest TCFL5 ChIP-seq peak to the transcription start site (TSS) for rhesus genes separated by class. Bigger circles indicate evolutionarily conserved piRNA genes. Vertical lines: median; whiskers: maximum and minimum values, excluding outliers. Values with the same individual values are indicated by a single marker indicating number of data points.

## Discussion

Our data, together with those from previous studies^5,11, 23, 30, 45^, suggest a model for the evolutionarily conserved transcriptional architecture by which A-MYB and TCFL5 collaborate to reprogram gene expression at the onset of male meiosis I (Fig. 7). Immediately before meiosis begins, a burst of retinoic acid induces transcription of *Stra8*. STRA8, in turn, activates *A-Myb* transcription during the leptotene or zygotene phases of meiotic prophase I, a period characterized by general transcriptional quiescence. By the onset of the pachynema, A-MYB initiates transcription of *Tcfl5*, and A-MYB and TCFL5 mutually reinforce their own transcription via interlocking positive feedback loops. Together A-MYB and TCFL5 directly promote transcription of hundreds of genes encoding proteins required for meiosis and spermiogenesis. These include components of the piRNA biogenesis machinery and the pachytene piRNA genes, whose transcripts are the sources piRNAs. Together and separately, A-MYB and TCFL5 participate in coherent feedforward loops that promote accumulation of piRNAs.

**Figure 7.**
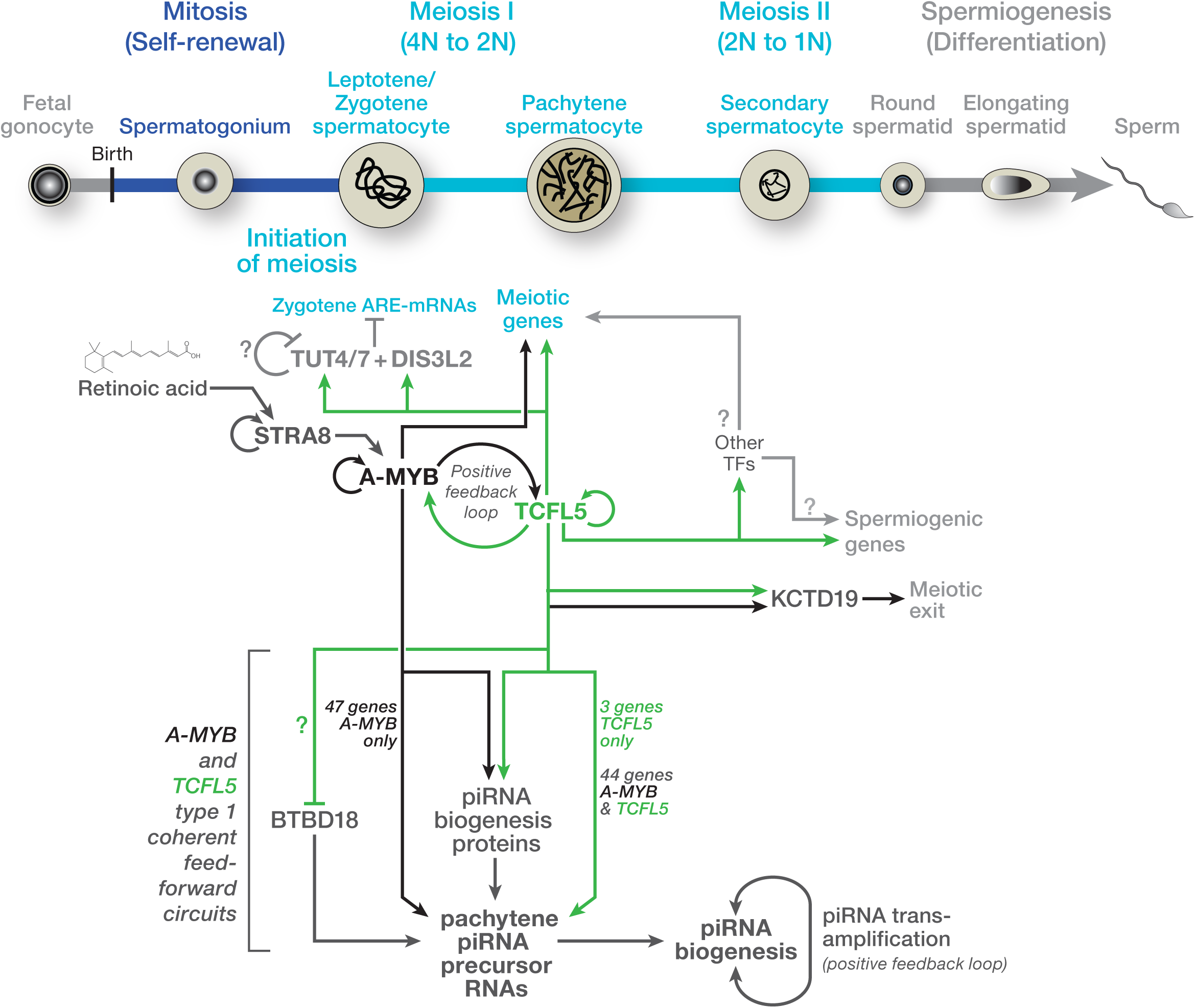
A model for the transcriptional architecture of mouse male meiotic cells. The model incorporates hypotheses from refs. 5,11,30,44,45 and this study. The developmental progression of spermatogenesis is aligned with the time of expression of various proteins. The figure highlights the proposed sequential roles of STRA8, A-MYB, and TCFL5, calling attention to the underlying architecture of the transcriptional circuits regulating male meiosis and piRNA production in mouse and rhesus macaque.

A-MYB and TCFL5 also promote transcription of additional transcription factors, chromatin-modifying enzymes, and at least one enzyme required for the biosynthesis of acyl-CoA, the acyl carrier used by the transferases that modify histone lysines to facilitate active transcription of many loci, including the pachytene piRNA genes. This transcriptional architecture is present in macaque, suggesting it is conserved among placental mammals. A-MYB also regulates male meiosis in chickens^11^. The chicken genome encodes TCFL5, raising the possibility that the overall architecture of the A-MYB/TCFL5 regulatory circuit predates the divergence of mammals and birds.

How do new pachytene piRNA-producing loci evolve? We suggest that three events promote the emergence of new pachytene piRNA loci: direct or indirect transcriptional activation of the locus by A-MYB or TCFL5; initiation of piRNA production from the newly transcribed locus by a piRNA from an older pachytene piRNA locus; and, finally, targeting of the same or another older pachytene piRNA gene by an initiator piRNA from the novel locus. Given that the majority of mouse pachytene piRNA genes are restricted to murine rodents or mice, cooption of non-coding transcripts into the piRNA pathway seems surprisingly easy. The resulting novel piRNAs are unlikely to provide a selectable advantage to the male mouse: surprisingly few piRNAs from even the most ancient pachytene piRNA genes appear to regulate genes required for the production of functional sperm^19, 24^. Supporting this view, human pachytene piRNA genes are among the most rapidly diverging sequences in human genome^12^. In fact, the main function of pachytene piRNAs seems to be to promote their own biogenesis. Do pachytene piRNAs really exist mainly to ensure their own production? We speculate that a small number of piRNA species modestly enhance spermatogenesis or spermiogenesis, thereby ensuring the inheritance of the tens of thousands of selfish piRNAs with no utility in germ cell development. Do most pachytene piRNAs benefit the mouse in subtle ways that remain to be discovered, or are pachytene piRNAs simply tiny narcissists that have turned the ancestral transposon silencing function of piRNAs against their host, joining the ranks of longer selfish genetic elements present in the mammalian genome? Distinguishing between these hypotheses promises to be both challenging and rewarding, like pachytene piRNAs themselves.

## Methods

### Mice

Mice were maintained and used according to the guidelines of the Institutional Animal Care and Use Committee of the University of Massachusetts Medical School (A201900331). C57BL/6J mice (RRID: IMSR_JAX:000664) were used as wild-type controls. *A-Myb^repro9^* (MGI3512907) were a gift from John Schimenti, and were maintained in a C57BL/6 background

To generate *Tcfl5^em1/em1^* mutant mice, we transcribed two sgRNAs targeting sequences in *Tcfl5* intron 1 (5′-GCA GUC UGG GUA CUA GAUA G-3′) and intron 3 (5′-AUU CAC UCA AAC AAC AAG AG-3′) using T7 RNA Polymerase. After purification by electrophoresis through a denaturing 10% polyacrylamide/urea gel, both sgRNAs (20 ng/µl each) and Cas9 mRNA (50 ng/µl, TriLink Biotechnologies, L-7206) were injected into the pronucleus and cytoplasm of fertilized eggs (Transgenic Animal Modeling Core, University of Massachusetts Medical School), and 15–25 blastocysts were transferred into uterus of pseudo-pregnant ICR females at day E2.5. Pups were screened by PCR of genomic DNA extracted from tail tissue using primers (5′-TGC CTC ATC TGG GTG GAT ATC TGA-3′; 5′-TAC ACG AAA ACA AAA CTT AAG CGG T-3′) designed to detect deletion of *Tcfl5* exons 2 and 3.

### Isolation of mouse germ cells by FACS

Germ cell sorting was as described^12, 13^. Briefly, testis was de-capsulated, incubated with 1× Gey′s Balanced Salt Solution (GBSS, Sigma, G9779) containing 0.4 mg/ml collagenase type 4 (Worthington; LS004188) at 33°C for 15 min, and the seminiferous tubules washed twice with 1× GBSS, and then incubated with 1× GBSS containing 0.5 mg/ml trypsin and 1 µg/ml DNase I at 33°C for 15 min. Next, the seminiferous tubules were maintained at 4°C on ice and gently pipetted up and down for 3 min through a Pasteur pipette to homogenize. Trypsin was inactivated with fetal bovine serum (FBS; f.c. 7.5% [v/v]), and the cell suspension was passed through a pre-wetted 70 µm cell strainer. Cells were recovered by centrifugation at 300 × *g* at 4°C for 10 min, resuspended in 1× GBSS containing 5% (v/v) FBS, 1 µg/ml DNase I, and 5 μg/ml Hoechst 33342 (Thermo Fisher, 62249), and incubated at 33°C for 45 min rotating at 150 rpm. Finally, propidium iodide (0.2 μg/ml, f.c.; Thermo Fisher, P3566) was added, and cells were filtered through a pre-wetted 40 µm cell strainer. Cell sorting (FACS Core, University of Massachusetts Medical School was as described^12, 53^.

### Testis histology and immunohistochemistry

Wild-type and *Tcfl5^em1/em1^* mutant testis tissue was fixed with Bouin’s solution, and paraffin embedded tissue sectioned at 5 μm thickness, and then stained with hematoxylin and eosin (H&E) solutions (Morphology Core Facility, University of Massachusetts Medical School). Immunohistochemistry (IHC) was performed using a standard protocol. Briefly, testis sections were de-paraffinized with xylene, dehydrated with ethanol, and then boiled in 1 mM citrate buffer (pH 6.0) to retrieve antigens. To inactivate endogenous peroxidase, tissue sections were incubated with 3% (v/v) hydrogen peroxide at room temperature for 10 min, then blocked with 5% (v/v) horse serum (ImmPRESS HRP Anti-Rabbit IgG [Peroxidase] Polymer Detection Kit; Vector labs; MP-7401). Sections were incubated with rabbit anti-TCFL5 (Sigma, HPA055223; 1:500 dilution), or rabbit anti-A-MYB (Sigma, HPA008791; 1:500 dilution) antibodies at 4°C overnight. Next, secondary HRP anti-rabbit antibody (Vector labs; MP-7401) was added and incubated at room temperature for 1 h, followed by incubation with chromogenic substrate (Fisher Scientific, TA-125-QHDX). Slides were counterstained with hematoxylin and covered with a #1 coverslip. IHC images were captured using a Leica DMi8microscope equipped with a 63×, 1.4 NA oil immersion objective.

### Sperm motility

Cauda epidydimal sperm were incubated in warm EmbryoMax HTF media at 37°C, 5% CO2. A drop of sperm was removed from the suspension and pipetted into a sperm counting glass chamber, then assayed by CASA or video acquisition. CASA was conducted using an IVOS II instrument (Hamilton Thorne, Beverly, MA) with the following settings: 100 frames acquired at 60 Hz; minimal contrast = 50; 4 pixel minimal cell size; minimal static contrast = 5; 0%straightness (STR) threshold; 10 μm/s VAP Cutoff; prog. min VAP, 20 μm/s; 10 μm/s VSL Cutoff; 5 pixel cell size; cell intensity = 90; static head size = 0.30–2.69; static head intensity = 0.10–1.75; static elongation = 10–94; slow cells motile = yes; 0.68 magnification; LED illumination intensity = 3000; IDENT illumination intensity = 3603; 37°C. The raw data files (i.e., .dbt files for motile sperm and .dbx files for static sperm) containing tracks ≥ 45 frames were used for sperm motility analysis. CASAnova was utilized to determine fractions of progressive and hyperactivated sperm from .dbt files of motile sperm, as previously described^54^

### Immunofluorescence

Immunofluorescent (IF) staining of FACS-purified germ cells was as described^13^. Briefly, freshly sorted germ cells were incubated in 25 mM sucrose solution at room temperature for 20 min and fixed with 1% (w/v) formaldehyde solution containing 0.15% (v/v) Triton X-100 at room temperature for 2 h. After washing the slides with (i) 1 × PBS containing 0.4% (v/v) Photo-Flo 200 (Kodak, 1464510), (ii) 1 × PBS containing 0.1% (v/v) Triton X-100, and (iii) 1 × PBS containing 0.3% (w/v) BSA, 1% (v/v) donkey serum (Sigma, D9663), and 0.05% (v/v) Triton X-100 (10 min each wash), slides were incubated with Rabbit polyclonal anti-SYCP3 (Abcam, ab15093; 1:1,000 dilution) and mouse monoclonal anti-γH2AX (Millipore, 05-636; 1:1,000 dilution) primary antibodies in 1 × PBS containing 3% (w/v) BSA, 10% (v/v) donkey serum, and 0.5% (v/v) Triton X-100 in a humidifying chamber at room temperature overnight. Next, slides were washed sequentially as (i–iii, above), then incubated with donkey anti-rabbit IgG (H+L) Alexa Flour 488 (Thermo Fisher Scientific, A-21206; 1:2,000) and anti-mouse IgG (H+L) Alexa Flour 594 (Thermo Fisher Scientific, A-21203; 1:2,000) secondary antibodies at room temperature for 1 h. The slides were washed three times with 1 × PBS containing 0.4% (v/v) Photo-Flo 200 (10 min each wash) and once with 0.4% (v/v) Photo-Flo 200 for 10 min.

For IF staining of cryopreserved testis sections, slides were fixed with 1 × PBS containing 0.1% (w/v) sodium periodate (NaIO_4_), 3% (w/v) L-lysine and 3% (v/v) paraformaldehyde at room temperature for 20 min. Slides were blocked with 2.5% goat serum. Next, slides were incubated with rabbit polyclonal anti-SYCP3 (Abcam, ab15093; 1:300 dilution) and mouse monoclonal anti-γH2AX (Millipore, 05-636; 1:300 dilution) antibodies at 4°C overnight. After primary antibody incubation, slides were washed twice with 1 × PBS containing 0.05% (v/v) Tween-20 and incubated with goat anti-rabbit IgG (H+L) Alexa Flour 568 (Thermo Fisher Scientific, A-11011; 1:500 dilution) and goat anti-mouse IgG (H+L) Alexa Flour 488 (Thermo Fisher Scientific, A-32723; 1:500 dilution) secondary antibodies at room temperature for 2 h. Next, nuclei were counterstained with 1 mg/ml DAPI (Thermo Fisher Scientific, 62248), the slides were sealed using gold antifade mountant solution (Fisher scientific, P36930) and covered with a #1 coverslip. IF images were captured using a Leica DMi8 Leica microscope equipped with a 63×, 1.4 NA oil immersion objective.

### RNA fluorescent in situ hybridization

Paraffin embedded testes tissue was sectioned at 5 μm thickness, then de-paraffinized with xylene and dehydrated with ethanol. RNAscope Hydrogen Peroxide (H_2_O_2_) (ACD, 322335) was used to inactivate endogenous peroxidase and antigens retrieved using RNAscope Target Retrieval Reagents (ACD, 322000) followed by treatment with RNAscope Protease Plus (ACD, 322331) according to the manufacturer’s directions. RNAscope 2.5 HD Detection Reagents-RED assay (ACD, 322360) were used to hybridize probes for *Tcfl5* (ACD, *Tcfl5* mRNA probe, NPR-0002511, targeting 202-1589 of NM_178254.3) and *A-MYB* (*A-MYB* mRNA probe, NPR-0002510, targeting 1165-2241 of NM_008651.3) mRNAs and to amplify and detect signal. Stained slides were sealed with EcoMount (Biocare Medical, EM897L) and imaged using a Leica DMi8 microscope equipped with a 63× 1.4 NA oil immersion objective. For simultaneous two-color detection of *Tcfl5* and *A-MYB* mRNAs, we used RNAscope two-color Fluorescent Detection v2 assay (ACD, 323110) and probes for *Tcfl5* (ACD, *Tcfl5* mRNA C3 probe, 815341-C3, targeting 202-1589 of NM_178254.3) and *A-MYB* (*A-MYB* mRNA C3 probe, 815331-C3, targeting 1165-2241 of NM_008651.3) mRNAs. Opal 520 reagent (Akoya Biosciences, FP1487001KT) was used to detect fluorescent signal for *Tcfl5* and Opal 690 reagent (Akoya Biosciences, FP1497001KT) for *A-Myb*. DAPI (1 mg/ml; Thermo Fisher Scientific, 62248) was used to counterstain nuclei. Stained slides were sealed using gold antifade mountant solution (Fisher scientific, P36930), covered with a #1 coverslip, and Z-stack images captured using a Leica SP8 Lightning confocal microscope equipped with a 63×, 1.4 NA oil immersion objective.

### TUNEL histochemical staining

ApopTag Plus Peroxidase In Situ Apoptosis Detection Kit (EMD Millipore; S7101) was used to detect DNA breaks according to the manufacturer’s protocol. In brief, Bouin solution-fixed, paraffin-embedded slides were de-paraffinized in three changes of xylene for 5 min each, gradually re-hydrated in 100% (v/v), 95% (v/v), and 70% (v/v) ethanol for 5 min each, and then washed in 1× PBS for 5 min. After pre-treating the slides with 20 µg/ml Proteinase K (EMD Millipore; 21627) at room temperature for 15 min, slides were washed with water twice (2 min each). Following pre-treatment, endogenous peroxidase was quenched in 3% (v/v) H_2_O_2_ at room temperature for 5 min. Slides were then incubated with equilibration buffer (EMD Millipore; Part # 90416) containing terminal deoxynucleotidyl transferase (EMD Millipore; Part # 90418) in a humidified chamber at 37°C for 1 h. Next, slides were incubated at room temperature for 10 min in 3% (v/v) stop/wash buffer (EMD Millipore; Part # 90419) to terminate the reaction and washed three times in 1× PBS for 5 min each. Slides were incubated with anti-digoxigenin conjugate (EMD Millipore; Part # 90420) in a humidified chamber at room temperature for 30 min, stained with peroxidase substrate at room temperature for 3 min. Nuclei were counterstained with hematoxylin, and the slides sealed with EcoMount (Biocare Medical, EM897L). Images were captured using Leica DMi8 microscope equipped with a 63×, 1.4 NA oil immersion objective.

### Western blotting

Frozen testis tissues were homogenized in a Dounce homogenizer using 20 strokes of pestle B in RIPA lysis buffer (25 mM Tris-HCl, pH 7.6, 150 mM NaCl, 1% (v/v) NP-40, 1% sodium deoxycholate, and 0.1% (w/v) SDS) containing 1× homemade protease inhibitor cocktail (1 mM 4-(2-Aminoethyl)benzenesulfonyl fluoride hydrochloride (Sigma; A8456), 0.3 μM Aprotinin, 40 μM Betanin hydrochloride, 10 μM. E-64 (Sigma; E3132), 10 μM Leupeptin hemisulfate). Lysed tissue was then sonicated (Branson Digital Sonifier; 450 Cell Disruptor) to break nuclei. After sonication, the samples were centrifuged at 20,000 × *g* at 4°C for 30 min. The supernatant was transferred to a new tube, and protein concentration measured using the Pierce BCA Protein Assay Kit (ThermoFisher; 23225). Total protein (50 µg) from each sample was mixed with 1/4 volume of loading dye (106 mM Tris-HCl, pH 6.8, 141 mM Tris base, 2% SDS, 10% v/v glycerol, 0.51 mM EDTA, 0.22 mM SERVA Blue G and 0.175 mM Phenol Red) containing 0.2 M dithiothreitol and heated at 95°C for 6 min to denature proteins. Samples were resolved by electrophoresis through a 4–20% polyacrylamide gradient SDS gel (Thermo Fisher, XP04205BOX), and the gel transferred to PVDF membrane (Millipore, IPVH00010). The membrane was blocked with Blocking Buffer (Rockland Immunochemicals, MB-070) at room temperature for 1.5 h and incubated with rabbit polyclonal anti-A-MYB (Sigma, HPA008791; 1:1,000 dilution), rabbit polyclonal anti-TCFL5 (Abgent, AP17006b; 1:1000 dilution; unless otherwise stated in the figure legends), or rabbit polyclonal anti-MILI (Abcam, ab36764; 1:1,000 dilution) antibody at 4°C overnight. Next, the membrane was washed three times (30 min each) with 1× PBS-T (0.1% (v/v) Tween-20 in 1× PBS) at room temperature; incubated with secondary donkey anti-rabbit IRDye 680RD antibody (LI-COR, 926-68073; 1:15,000 dilution) at room temperature for 30 min; washed three times (30 min each) with 1× PBS-T at room temperature, and signal detected using the Odyssey Infrared Imaging System (LI-COR). As a loading control, membrane was incubated with mouse anti-ACTIN antibody (Santa Cruz Biotechnology, sc-47778; 1:10,000 dilution) at room temperature for 2 h, followed by goat anti-mouse IRDye 800RD secondary antibody (LI-COR, 926-32210; 1:15,000 dilution) as described above.

### Small-RNA library construction and analysis

Total RNA was extracted from frozen testis tissue using the mirVana miRNA isolation kit (Thermo Fisher, AM1560). Small RNA library construction was performed as described^13^. Briefly, total RNA was resolved by electrophoresis through a denaturing 15% polyacrylamide gel to select 18–34 nt RNAs. Size-selected small RNA was oxidized with sodium periodate (NaIO_4_) prior to 3′ adaptor ligation to suppress ligation to microRNAs and siRNAs. The 3′ adaptor (5′-rApp NNN TGG AAT TCT CGG GTG CCA AGG /ddC/-3′) was ligated in the presence of 50% (w/v) PEG8000 (Rigaku, 1008063). The 3′ adaptor ligated small RNAs were then purified by electrophoresis through a denaturing 15% polyacrylamide gel to eliminate the unligated 3′ adaptor; 42 to 60 nt small RNA containing the 3′ adaptor were recovered from the gel. Finally, 5′ adaptor was ligated to the RNA and cDNA generated. Libraries were sequenced (79 nt single-end reads) using a NextSeq500 (Illumina).

After removing 3′ adaptor sequence from the reads, we eliminated reads with PHRED score < 5. Remaining reads were then mapped to mouse genome (mm10) using piPipes^18^ employing Bowtie 2.2.5 (ref. 55), allowing one mismatch. We quantified the length distribution and abundance of piRNA reads by normalizing to genome-mapping reads reported as reads per million (rpm). Small RNA sequencing statistics are provided in Supplementary Table 7.

### Long RNA library construction and analysis

Total RNA from frozen testis samples and from sorted germ cells was extracted using mirVana miRNA isolation kit (Thermo Fisher, AM1560). Before RNA library construction from FACS-purified germ cells, we added 1 µl of 1:100 dilution of ERCC spike-in mix 1 (Thermo Fisher, 4456740, LOT00418382) to 0.5–1µg total RNA to enable subsequent molecular quantification of transcripts. Library construction was performed as described^56^ with a modified ribosomal RNA (rRNA) removal procedure. Briefly, 186 short antisense DNA oligos (ASOs; 50–80 nt) complementary to mouse rRNAs were designed and pooled (0.05 µM/each)^57, 58^. Pooled rRNA ASO (1 µl) were added to 1 µg total RNA in the presence of 100 mM Tris-HCl (pH 7.4) and 200 mM NaCl, heated to 95°C and cooled to 22°C at −0.1°C/s then incubated at 22°C for 5 min. The DNA:RNA hybrids were digested with 10 U Thermostable RNase H (Epicentre, H39500) in 50 mM Tris-HCl (pH 7.4), 0.1 M NaCl, and 20 mM MgCl_2_ at 45°C for 30 min. Next, samples were treated with 4 U Turbo DNase (Thermo Fisher, AM2238) at 37°C for 20 min. To enrich RNA >200 nt and remove tRNA, RNA samples were purified with an RNA Clean & Concentrator-5 (Zymo Research, R1015), and cDNA generated as described. Libraries were sequenced as 79 + 79 nt paired-end reads using a NextSeq500 (Illumina).

rRNA reads were eliminated using Bowtie 2.2.5 with default parameters^55^, and then the remaining reads were mapped to the mouse genome (mm10) using STAR 2.3 (ref. 59). Results in SAM format were de-duplicated and transformed into BAM format using SAMtools 1.8 and Umitools^60, 61^. Finally, mapped reads were assigned to protein-coding genes, long non-coding RNAs (lncRNAs) and piRNA genes using HTSeq 0.9.1 (ref. 62). Transcript abundance was reported as reads per million uniquely mapped reads per thousand nucleotides (RPKM) calculated using homemade Bash scripts.

Transcript abundance in molecules per cell was performed as described^13^. Because each sample contained ∼623,291,645 molecules of ERCC spike-in mix, the abundance of each gene = (number of mapped reads × 623291645)/(number of cells used to prepare the library × the number of reads mapping to the ERCC spike-in sequences). RNA sequencing statistics are provided in Supplementary Table 7. All analyses used Ensembl-v86 gene annotations.

### Chromatin immunoprecipitation and sequencing

Chromatin Immunoprecipitation (ChIP) was performed as described^11, 12^. Briefly, frozen testis tissue was minced on dry ice and incubated in 1.7 ml tubes in ice-cold PBS containing 2% (w/v) formaldehyde at room temperature for 30 min, slowly tumbling end-over-end. Fixed tissue was crushed in ChIP lysis buffer (1% SDS, 10 mM EDTA, 50 mM Tris-HCl, pH 8.1) with a Dounce homogenizer using 40 strokes pestle B (Kimble-Chase, Vineland, USA). To shear chromatin to 150–200 bp, samples were sonicated (Covaris, E220). Sonicated lysate was then diluted 1:10 with ChIP dilution buffer (0.01% SDS, 1.1% Triton X-100, 1.2 mM EDTA, 16.7 mM Tris-HCl, pH 8.1, 167 mM NaCl) and immunoprecipitated using 5 µg anti-TCFL5 (Sigma, HPA055223, lot # 000000907) or 5.5 µg anti-A-MYB (Sigma, HPA008791, lot # C105987) antibody. After immunoprecipitation, DNA was extracted with phenol:chloroform:isoamyl alcohol (25:24:1; pH 8), and ChIP-seq libraries generated as described^56^. Libraries were sequenced as 79 + 79 nt paired-end reads using a NextSeq500 (Illumina).

Raw reads were mapped to the mouse genome (mm10) using Bowtie 2.2.5 with parameters `—very-sensitive —no-unal —no-mixed —no-discordant -|10`. Duplicate reads were removed using SAMtools 1.8 (ref. 61). Normalized genome coverage files in bigWig format were generated using a homemade script. Data are reported as reads per million (rpm) using uniquely mapping reads normalized to read depth. Significant TCFL5 or A-MYB peaks (FDR < 0.05 for enrichment relative to input) were identified using MACS 1.1.2 (ref. 63) with parameters `-keep-dup all –q 0.001`. ChIP sequencing statistics are provided in Supplementary Table 7.

The distance (bp) of transcription start site to the nearest TCFL5 or A-MYB peak summit for each gene was calculated. Genes that retain TCFL5 or A-MYB peak within the range of 500 bp around their transcription start site were considered as TCFL5- or A-MYB-regulated genes.

### CUT&RUN Sequencing

Cleavage Under Targets and Release Using Nuclease (CUT&RUN) sequencing from FACS-purified germ cells was performed as described^29^. Briefly, FACS-purified germ cells were lysed using nuclear extraction buffer (20 mM HEPES-KOH, pH 7.9, 10 mM KCl, 0.5 mM Spermidine, 0.1% Triton X-100, 20% glycerol) and nuclei recovered by centrifugation at 600 × *g* at 4°C for 3 min. Nuclei were immobilized on BioMag Plus Concanavalin A-coated paramagnetic beads (Bangs Laboratories, BP531) at room temperature for 20 min. After immobilizing on beads, nuclei were blocked in 20 mM HEPES-KOH, pH 7.5, 150 mM NaCl, 0.5 mM spermidine, 0.1% BSA, 2 mM EDTA) at room temperature for 5 min. Then 2.5 µg anti-TCFL5 (Sigma, HPA055223 or Sigma, HPA055223, lot # 000000907) or 2.5 µg anti-A-MYB (Sigma, HPA008791, lot # C105987) antibodies were added to the beads and incubated at 4°C for 4 h, slowly tumbling end-over-end. CUT&RUN without antibody provided a negative control. After antibody incubation, beads were washed twice in wash buffer (20 mM HEPES-KOH, pH 7.5, 150 mM NaCl, 0.5 mM spermidine, 0.1% BSA) and incubated with 1.5 µl protein A-MNase fusion protein (pA-MN; 140 µg/ml) at 4°C for 2 h, slowly tumbling end-over-end. To remove unbound pA-MN, beads were washed twice in wash buffer. To liberate DNA bound by TCFL5 or A-MYB, 3 µl 100mM CaCl_2_ was added to the beads and incubated at 0°C for 7 min. The reaction was stopped by adding an equal volume of 2× STOP buffer (200 mM NaCl, 20 mM EDTA, 4 mM EGTA, 50 µg/ml RNase A, 40 µg/ml glycogen) and then incubated at 37°C for 20 min. Nuclei were removed by centrifugation at 16,000 × *g* at 4°C. The supernatant containing CUT&RUN fragments was transferred to a fresh 1.7 ml tube. After extracting the DNA with phenol:chloroform:isoamyl alcohol (25:24:1; pH 8), CUT&RUN libraries were prepared as described^56^ with slight modifications. Briefly, end-repair using 0.5 U/µl T4 DNA polymerase (NEB, M0203) and adenylation using 0.5 U/µl *Taq* DNA polymerase (NEB, M0273) were carried out in a single reaction. Then, end-repaired and adenylated DNA was purified using AMPure beads. Finally, Illumina adapters (MP-Ada1: 5′-p GAT CGG AAG AGC ACA CGT CT-3′; MP-Ada2: 5′-p ACA CTC TTT CCC TAC ACG ACG CTC TTC CGA TCT-3′) were added using 600 U/µl T4 DNA ligase (Enzymatics Inc., L603-HC-L) at room temperature for 30 min.

CUT&RUN fragments were sequenced using 79 + 79 nt paired-end reads on a NextSeq500 (Illumina). Raw reads were mapped to the mouse genome (mm10) using Bowtie 2.2.5 (ref. 55) with parameters `—very-sensitive —no-unal —no-mixed — no-discordant -|10`. TCFL5 and A-MYB peaks were identified used <u>S</u>parse <u>E</u>nrichment <u>A</u>nalysis for <u>C</u>UT&<u>R</u>UN (SEARC^43^) in “norm relaxed” mode. CUT&RUN performed without antibody was used as the background control. Genes with a TCFL5 or A-MYB peak ±500 bp of their transcription start site were considered as TCFL5- or A-MYB-regulated. To be considered bound by TCFL5, a gene was required (1) to have a SEARC-identified peak within 500 bp of its TSS in both CUT&RUN replicates; or (2) to have a SEARC-identified peak within 500 bp of its TSS in one of the two CUT&RUN replicates and a MACS2-identified peak within 500 bp of its TSS in two of three ChIP-seq replicates. To be considered bound by A-MYB, a gene was required (1) to have a SEARC-identified peak within 500 bp of its TSS in both CUT&RUN replicates; or (2) to have a SEARC-identified peak within 500 bp of its TSS in one of the two CUT&RUN replicates and a MACS2-identified peak within 500 bp of its TSS in ChIP-seq. CUT&RUN sequencing statistics are provided in Supplementary Table 7.

### Statistics

All statistics were calculated using R console (https://www.rstudio.com); graphs were generated using R console or Prism 8.4.3 (GraphPad Software, LLC.). For box plots, boxes represent the first and third quartiles; except in Fig. 3, whiskers show the maximum and minimum values excluding outliers. Outliers are defined as data points with values > third quartile + 1.5 × IQR or lower than the first quartile − 1.5 × IQR, where IQR (interquartile range) is the maximum third quartile minus the first quartile minimum. Two-sided Mann-Whitney-Wilcoxon U test was used to calculate *p* values.

## Author contributions

D.M.Ö., and P.D.Z. conceived and designed the experiments. D.M.Ö., K.C., H.M., A.A., C.C., A.B. and I.G. performed the experiments. Y.T.X., and D.M. Ö. analyzed the sequencing data. D.C.P., and D.G.R. characterized the *Tcfl5^em1/em1^* testis phenotype. D.M.Ö and P.D.Z. wrote the manuscript. All authors discussed and approved the manuscript.

## Data Availability

Sequencing data are available from the National Center for Biotechnology Information Sequence Read Archive using Gene Expression Omnibus accession number GSE166104.

## Code availability

Code used to identify putative piRNA-directed cleavage sites is available at https://github.com/weng-lab/GTBuster.

## Acknowledgements

We thank S. Pechold, T. Giehl, B. Gosselin, Y. Gu and T. Krumpoch at UMMS FACS Core for their expert mouse germ cell sorting; J. Gallant at the UMMS Transgenic Animal Modeling Core for help generating *Tcfl5^em1/em1^* mutant mice; K. Orwig for providing rhesus macaque testis specimens; C. Baer for her expert help with Leica SP8 Lightning confocal microscopy; C. Tipping and C. Andersson for technical assistance; and members of the Zamore laboratory for discussions and critical comments on the manuscript.

---

**Extended Data Figure 1.**
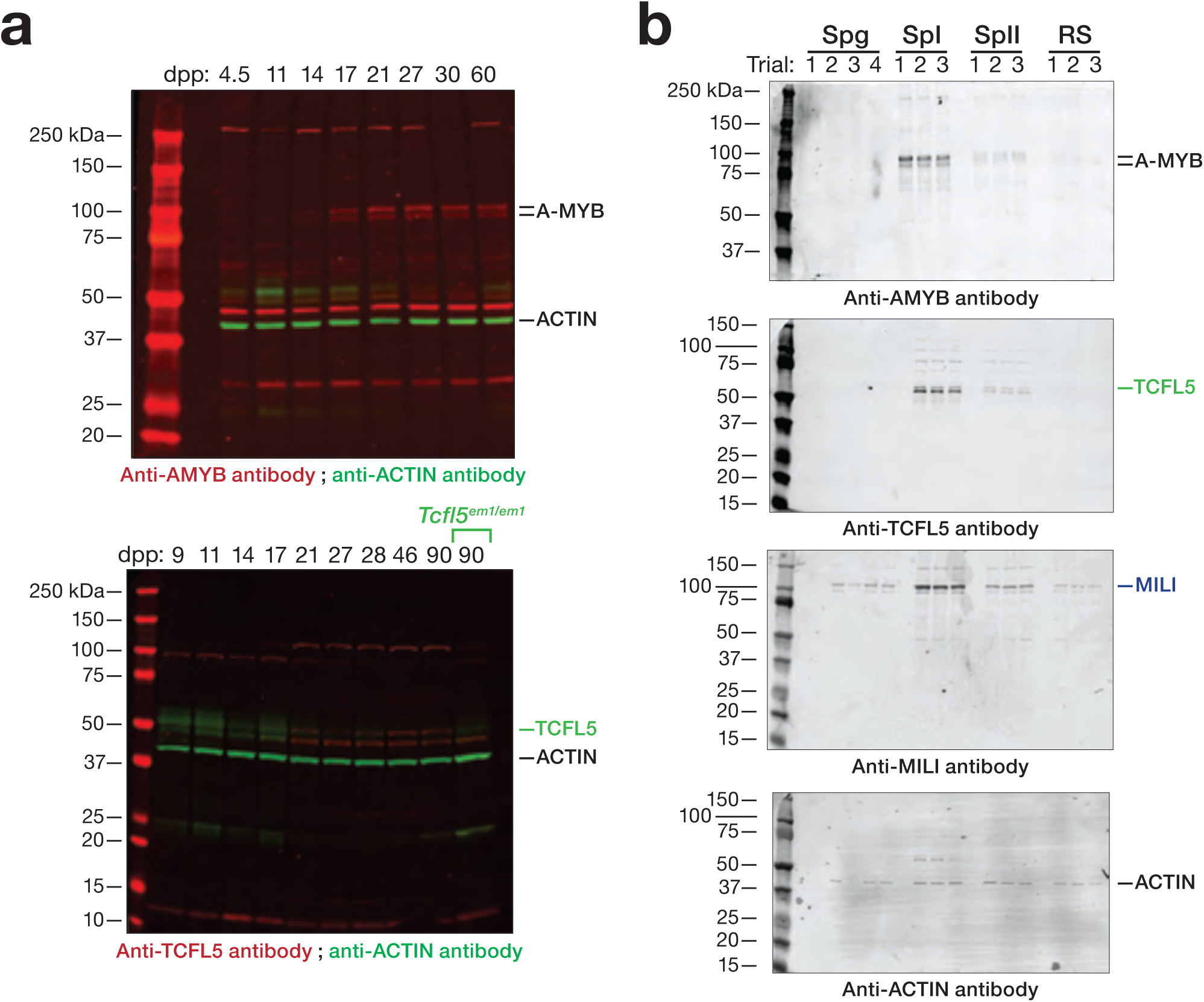
Temporal and spatial expression of A-MYB and TCFL5. **a, b,** Protein abundance of A-MYB and TCFL5 from (**a**) staged mouse testis (50 µg protein of testis lysate per lane) or (**b**) FACS-purified germ cells (protein lysate from ∼100,000 germ cells per lane). ACTIN served as a loading control.

**Extended Data Figure 2.**
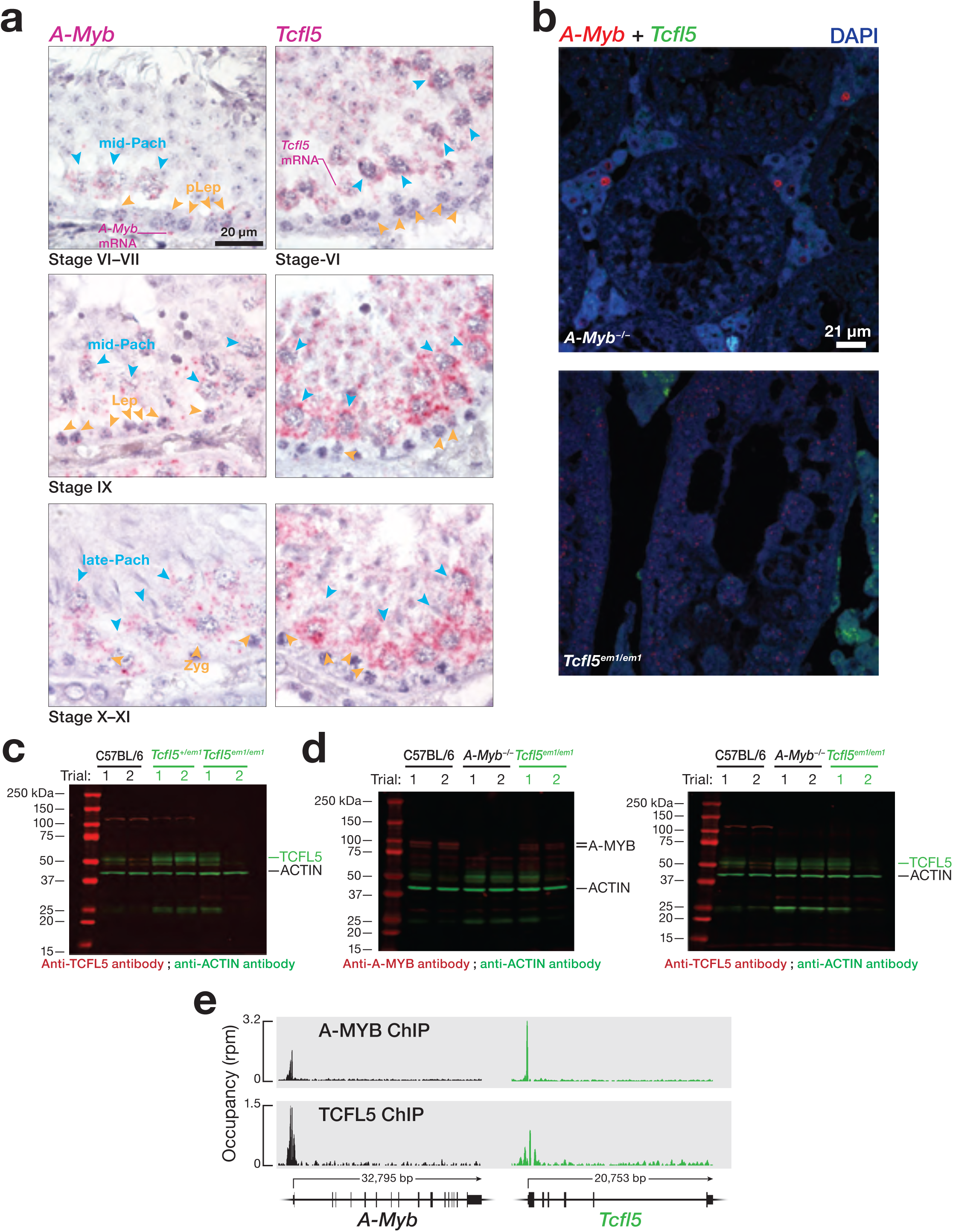
Reciprocal positive feedback loops between A-MYB and TCFL5. **a**, Temporal mRNA expression of *A-Myb* and *Tcfl5* for a seminiferous tubule section of C57BL/6 mouse adult testis detected by RNA fluorescent *in situ* hybridization. **b,** *A-Myb* and *Tcfl5* mRNAs were detected in the seminiferous tubules of *Tcfl5^em1/em1^* and *A-Myb^−/−^* mutant mice using two-color RNA fluorescent *in situ* hybridization. **c,** Abundance of TCFL5 protein in *Tcfl5^em1/em1^* and *Tcfl5^+/em1^* mutant testes, compared to C57BL/6 wild-type was measured by immunoblotting. ACTIN serves as a loading control. Each lane contained 50 µg protein of testis lysate. **d,** Protein abundance of A-MYB and TCFL5 from *A-Myb^−/−^* and *Tcfl5^em1/em1^* mutant mice testes was measured by immunoblotting. ACTIN serves as a loading control. Each lane contained 50 µg protein of testis lysate. **e,** A-MYB and TCFL5 occupancy at the promoters of the *A-Myb* and *Tcfl5* genes was measured using ChIP-seq.

**Extended Data Figure 3.**
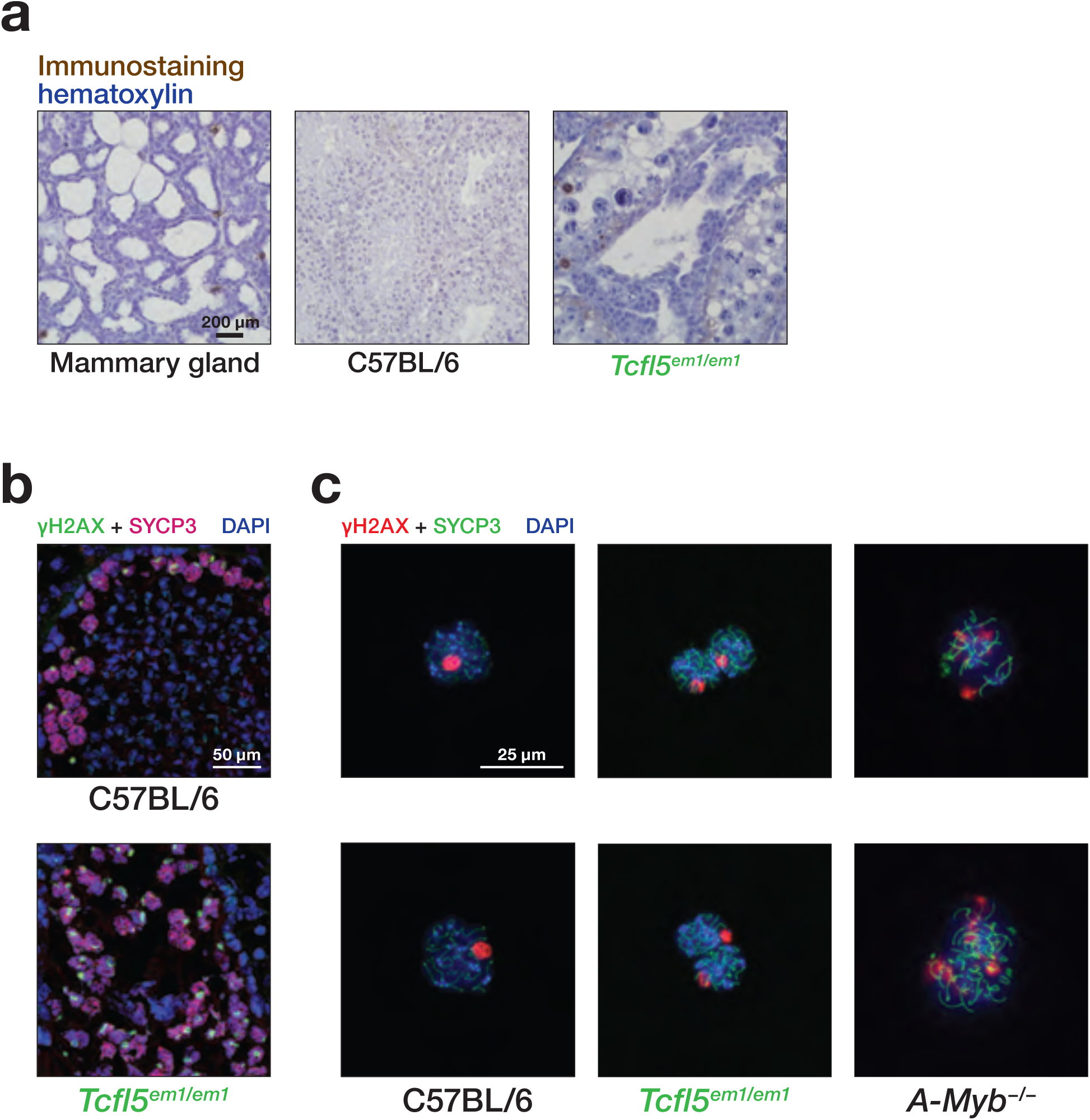
DNA double strand breaks (DBSs) are repaired in *Tcfl5^em1/em1^* but not *A-Myb^−/−^* mice. **a,** Terminal deoxynucleotide transferase dUTP nick end-labeling (TUNEL) assay was used to detect apoptotic cells. Female mammary gland serves as positive control. **b,** Immunofluorescent staining was used to detect Synaptonemal Complex Protein 3 (SYCP3) and γH2AX proteins in cryopreserved C57BL/6 and *Tcfl5^em1/em1^* mouse testes sections. **c,** Immunostaining to detect SYCP3 and γH2AX proteins in meiotic spreads of spermatocyte nuclei. DAPI was used to mark double-stranded DNA

**Extended Data Figure 4.**
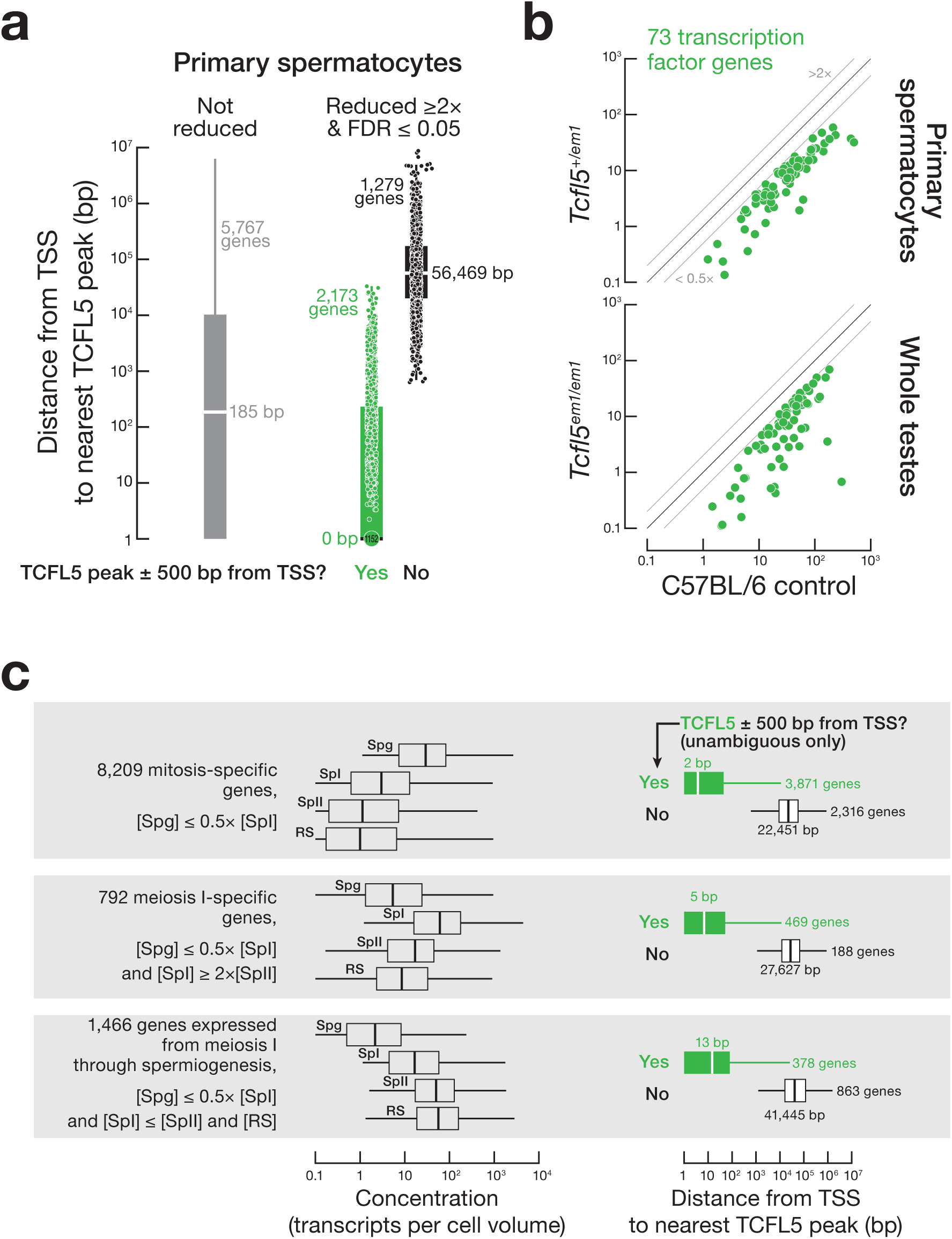
TCFL5 regulates genes required for meiosis and spermiogenesis and genes encoding downstream transcription factors. **a**, Boxplots display the mean distance of two replicates from the annotated transcription start site (TSS) to the nearest TCFL5 peak determined by CUT&RUN for genes whose transcripts were significantly reduced in primary and secondary spermatocytes and round spermatids purified by FACS from *Tcfl5^+/em1^* testes. Genes classified as not bound by TCFL5 showed no TCFL5 peak ±500 bp from the TSS in any CUT&RUN or ChIP-seq datasets. Vertical lines: median. Whiskers: maximum and minimum values, excluding outliers (i.e., 1.5 × IQR). Measurements with the same values are indicated by a single marker indicating number of individual data points. **b**, Scatter plot of the steady-state mRNA abundance for 73 TCFL5-bound, transcription factor-encoding genes in *Tcfl5^+/em1^* primary spermatocytes or *Tcfl5^em1/em1^* whole testes compared to C57BL/6 controls. Data points correspond to the mean of three C57BL/6 and three *Tcfl5^+1/em1^* trials (top panel) or six C57BL/6 and two *Tcfl5^em1/em1^* trials. **c**, Box plots show transcript concentration (i.e., abundance normalized to cell volume) for three classes of genes in spermatogonia (Spg), primary (SpI) and secondary (SpII) spermatocytes, and round spermatids (RS). Vertical lines: median; whiskers: maximum and minimum values, excluding outliers (i.e., 1.5 × IQR). **d**, The distance (mean of three replicates) from the nearest TCFL5 peak, detected by ChIP-seq, to the transcription start site (TSS) for three classes of genes. Genes were only scored as not bound by TCFL5 if they had no TCFL5 peak ±500 bp from the TSS in any CUT&RUN or ChIP-seq datasets. Vertical lines: median. Whiskers: maximum and minimum values, excluding outliers (i.e., 1.5 × IQR).

**Extended Data Figure 5.**
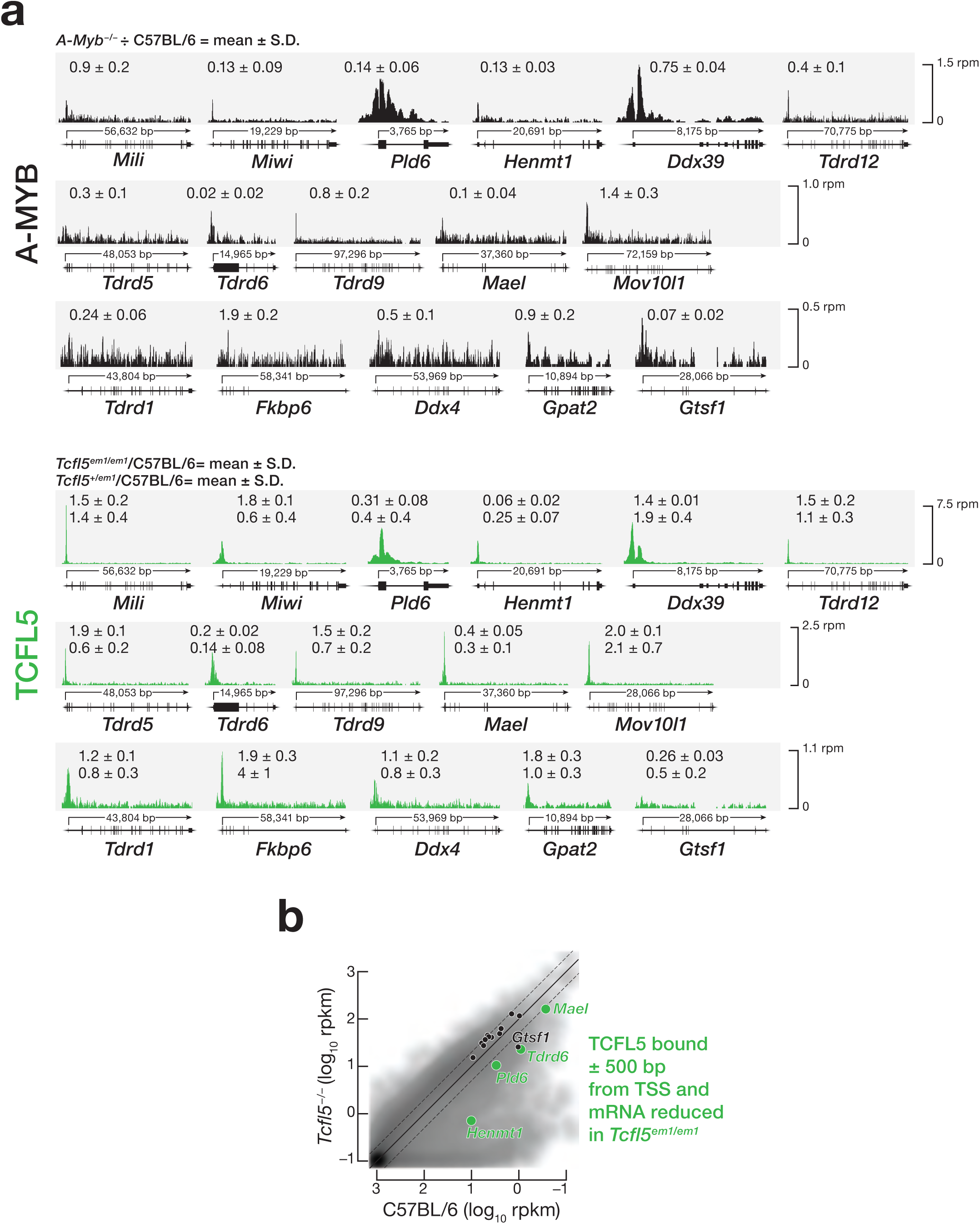
TCFL5 drives the transcription of mRNAs encoding mouse piRNA biogenesis proteins. **a,** A-MYB and TCFL5 CUT&RUN peaks at the promoters of 16 genes that encode piRNA biogenesis proteins. **b,** Scatter plot of the steady-state mRNA abundance of piRNA biogenesis genes in *Tcfl5^em1/em1^* mutant mice.

**Extended Data Figure 6.**
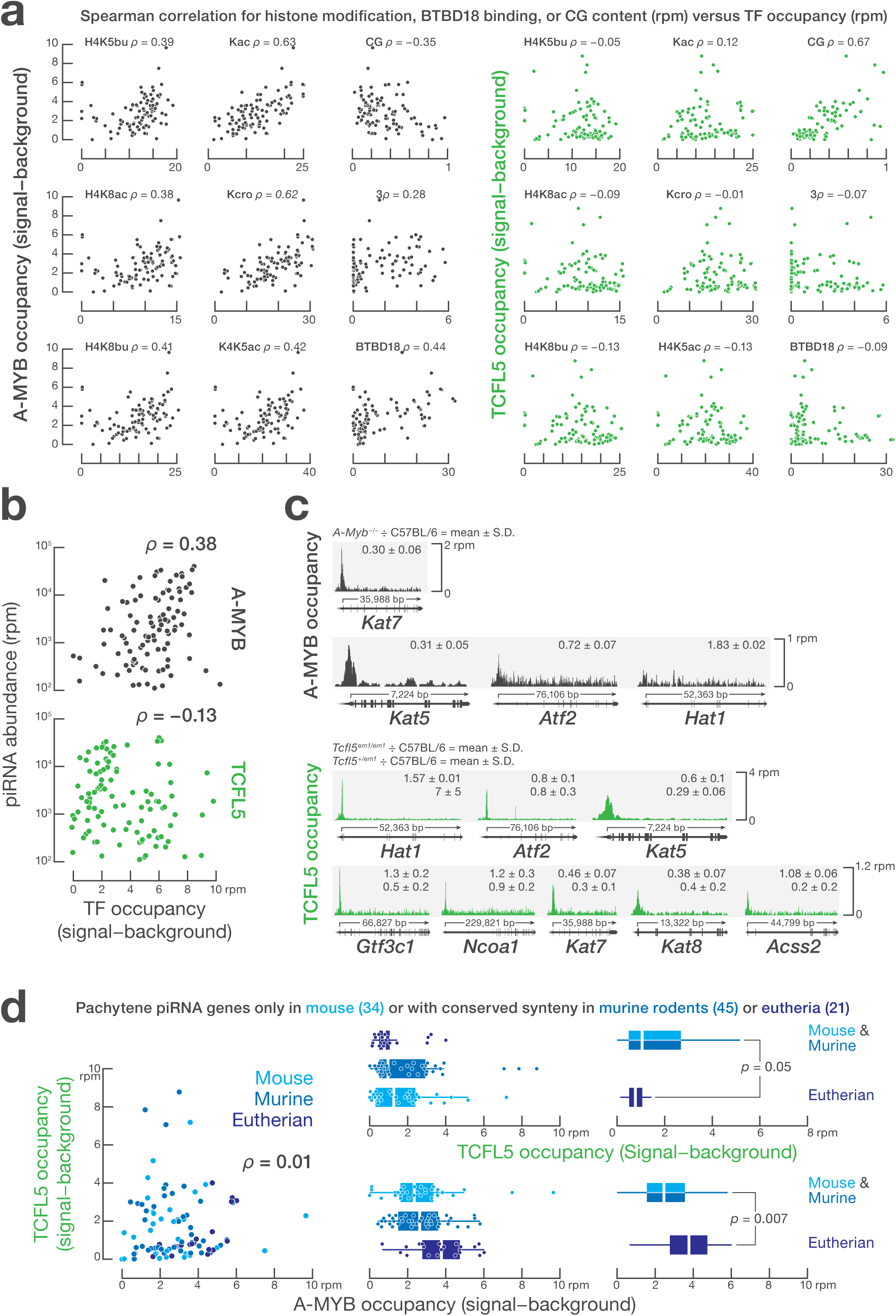
TCFL5 regulates evolutionarily younger, pachytene piRNA-producing genes. **a,** Spearman correlation (ρ) between A-MYB or TCFL5 occupancy and the evolutionarily conserved features of pachytene piRNA genes. **b,** A-MYB and TCFL5 CUT&RUN peaks at the promoters of genes encoding histone acylation enzymes: *Hat1*, *Atf2*, *Kat5*, *Kat6a*, *Gtf3c1*, *Ncoa1*, *Kat7*, *Kat8*, and *Acss2*. The change in steady-state mRNA expression in *Tcfl5^em1/em1^* whole testes and *Tcfl5^+/em1^* mutant primary spermatocytes, relative to C57BL/6 controls, is reported for each gene as mean ± SD. **c**, Spearman correlation between A-MYB or TCFL5 occupancy and piRNA abundance. **d**, Left panel, scatter plot to test the Spearman correlation between A-MYB and TCFL5 occupancy around the TSS of pachytene piRNA genes classified by conservation of their genomic locations (synteny) among eight placental mammals. Center panels, A-MYB and TCFL5 occupancy, determined using CUT&RUN, at the promoters of the same three class of pachytene piRNA genes. Vertical lines: median; whiskers: maximum and minimum values excluding outliers (i.e., 1.5 × IQR). Each dot represents an individual pachytene piRNA gene. Right panels, same as in the center panels, except comparing A-MYB and TCFL5 occupancy for the promoters of pachytene piRNA genes with or without conserved synteny among eutherian mammals. *p*-value was calculated using a two-sided Mann-Whitney-Wilcoxon U test.

